# SALMONID SCIENCE ON THE MOVE: TWENTY YEARS OF RESEARCH THROUGH THE LENS OF EARLY CAREER RESEARCHERS

**DOI:** 10.64898/2026.10.26.761798

**Authors:** A. Fontaine, L. E. Hunter, A. Lamarins, W. B. Perry, M. Roncoroni

## Abstract

Salmonids are among the most studied fish species worldwide, owing to their wide geographic distribution, diversity of life-history strategies, ecological and socioeconomic importance, and vulnerability to global change. In this review, we take advantage of two decades of the *International Workshop of PhDs and Post-doctoral Fellows on Anadromous Salmonids* (NoWPaS) conference series to track the evolution of salmonid science through the lens of early-career researchers (ECRs). From all abstracts presented at NoWPaS meetings between 2005 and 2025, we extracted structured qualitative and quantitative information to characterize major trends and shifts in the field. Our results show a strong research bias toward Atlantic salmon and brown trout, concentrated in Europe and Atlantic Canada, and a persistent mismatch between the life stages and habitats studied. In particular, the marine phase remains understudied in comparison to fresh and coastal waters. Over the study period, salmonid research shifted markedly from hydro-geomorphological and habitat-based approaches toward genetic and genomic methods, alongside the rise of advanced telemetry methods for reconstructing individual life histories. Research orientation, measured as a gradient from fundamental to applied research, was on average intermediate but varied among countries and disciplines, with the UK and Norway showing shifts to more applied research, as well as the discipline of *Ecology* becoming more applied. Additionally, we identified a significant increase in *Climate change* and *Aquaculture* as research drivers, with some drivers well-represented in some species (*e.g*., *Hydropower and dams* for landlocked Atlantic salmon and *Climate change* for Arctic char). These trends should be interpreted in context of the composition of the NoWPaS network: participation is primarily ECRs in the Northern Hemisphere, with research from Asia, South America, and western North America largely absent from this synthesis. Despite this lens, our analysis identifies clear knowledge gaps, particularly around marine life stages and species other than Atlantic salmon and brown trout, and highlights opportunities for broadening taxonomic, geographic, and life-history coverage in future salmonid research. We also present a roadmap, identifying combinations of life stages, disciplines and taxa which have been under-represented and constitute potential knowledge gaps, while also flagging those combinations that have already been well studied. Overall, we advocate for a continued shift toward integrative and holistic approaches, bridging distinct research disciplines, to better understand, manage, and conserve salmonids under increasingly uncertain future environmental conditions.

**GRAPHICAL ABSTRACT:** 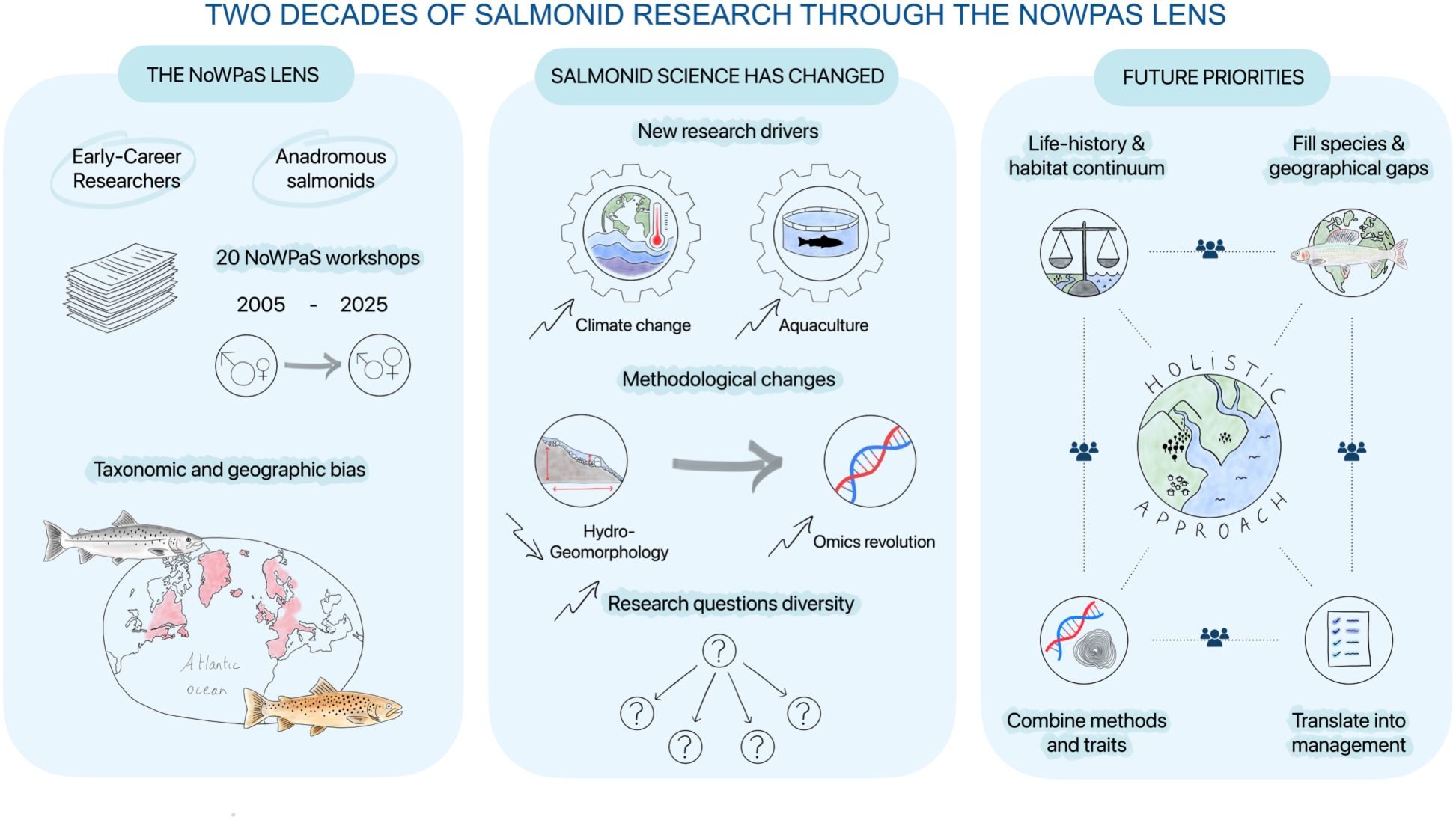

## 1 INTRODUCTION

Salmonids are cold-water adapted fish species native to the Northern Hemisphere and introduced in many temperate or cold regions (*e.g.*, Chilean Patagonia) of the Southern Hemisphere. They show remarkable life-histories throughout their native and introduced ranges, including full freshwater residency (*e.g.*, grayling, *Thymallus* spp.; Englmaier et al., 2022) or (partial) anadromy (*e.g*., brown trout, *Salmo trutta*; Nevoux et al., 2019), iteroparity (*e.g*., Arctic char, *Salvelinus alpinus*; Birnie-Gauvin et al., 2023) or semelparity (*e.g*., Pacific salmon, *Oncorhynchus* spp.; Crespi and Teo, 2002). Even within the same species, multiple life strategies can be observed from complete freshwater residency to long-distance migratory behaviors (Birnie-Gauvin et al., 2019), making salmonids a well-established illustration of ecological biocomplexity in natural systems. Salmonids are also recognized for their ecological, cultural, recreational, and economic importance. For instance, the decomposition of Pacific salmon carcasses promotes the transfer of nutrients from the sea to both terrestrial and freshwater ecosystems (Schindler et al., 2003; Moore et al., 2007; Walsh et al., 2020), Arctic char are central to the diet and culture *(e.g*., community gatherings, feasts) of many Indigenous communities across the coastal Canadian Arctic (Bolduc et al., 2025) and Atlantic salmon (*Salmo salar*) made up 15.5% of all global marine aquaculture production by value in 2023 (FAO, 2025).

Nevertheless, salmonid populations are reported as being at risk and/or in decline across the Northern Hemisphere, for both anadromous (*e.g*., Crozier et al., 2021; Dadswell et al., 2022; Svenning et al., 2024) and non-anadromous (*e.g*., Burkhardt-Holm et al., 2002; Vom Berge et al., 2025) populations. For example, worldwide nominal catches of Atlantic salmon collapsed by around 80% between 1975 and 2020 (ICES, 2021). The causes of salmonids’ decline are multiple and often interrelated, especially for species with complex life cycles that are continuously exposed to a variety of external stressors. For instance, the decline of anadromous populations has been attributed to higher mortality in the marine phase due to climate driven changes in oceanic conditions, increased predation and overfishing, but also losses in freshwater due to stressors such as parasites and diseases (*e.g*., proliferative kidney disease), invasive species, climate driven changes in hydrology, habitat modification and changes to water quality; this is not to mention the loss of genetic diversity (which underpins resilience to environmental change) due to fish farming escapees and stocking (*e.g.*, Bradford and Irvine, 2000; Borsuk et al., 2006; Lobón-Cerviá, 2009; Williams et al., 2009; Almodóvar et al., 2012; Al-Chokhachy et al., 2016; Olmos et al., 2020; Bell et al., 2021; Flávio et al., 2021; Shephard and Gargan, 2021; Tonina et al., 2022; Almodóvar et al., 2023; van Rijssel et al., 2024; Tissot et al., 2025). These same freshwater stressors are also causing declines in non-anadromous populations. Yet, disentangling how stressors influence salmonids in a time of global change remains particularly challenging due to their complex life histories, interactions between stressors, as well as the different levels in which stressors act, from individuals to populations (Crozier and Siegel, 2023).

Owing to their broad geographic distribution, biocomplexity, ecological and socio-economic importance, and the severity of their current decline, salmonids have become model organisms for addressing a wide range of ecological, evolutionary, and conservation questions, as illustrated by several recent reviews, syntheses, and research papers (*e.g.,* Garcia de Leaniz et al., 2007; Jonsson and Jonsson, 2009; Keefer and Caudill, 2014; Birnie-Gauvin et al., 2019, 2021, 2023; Pitman et al., 2021; Munsch et al., 2022; Crozier and Siegel, 2023; Lamarins et al., 2024; Roncoroni, 2024, Fontaine et al. 2025). This expanding body of work is produced by a large and diverse research community, often spanning freshwater, estuarine, and marine systems, as well as multiple salmonid species and multiple disciplines, from geneticists to hydrologists. Given the accelerating impacts of global change on salmonids, collaborative research is required to enhance adaptive capacity and support evidence-based decision-making by managers and policymakers toward a common conservation objective. In this context, networks that facilitate exchange, collaboration, and synthesis, particularly among early-career researchers (ECRs), are essential to ensure a coherent development and integration of research efforts across institutions and countries (Betts et al., 2026a).

In response to this growing and diverse research community, the *International Workshop of PhDs and Post-doctoral Fellows on Anadromous Salmonids*, referred to as NoWPaS, was established as an annual scientific workshop organized by and for research-oriented master students, doctoral and post-doctoral researchers working on salmonid research. The NoWPaS network was established in 2005 by Norwegian, Swedish, Finnish and Danish researchers with the primary goal of fostering interdisciplinary discussions and collaborations among ECRs to shape and lead current and future research on salmonids, with each meeting also hosting keynote presentations from senior researchers leading in their field. Since then, the network has grown internationally, expanding beyond the Nordic community, with workshops held in Norway (2005, 2012, 2023), Sweden (2006, 2014), Finland (2007, 2018), Denmark (2008), the United Kingdom (2009, 2010, 2013, 2017, 2019, 2024), France (2011, 2022), Ireland (2015), Canada (2016, 2025), Iceland (2020, 2026), and once virtually (2021). Up to 2025, NoWPaS workshops hosted 633 presentations, 88% of which were from master students, doctoral and postdoctoral researchers.

Building on this significant contribution, this review synthesizes the abstracts presented at NoWPaS workshops to provide an overview of past and present salmonid research. The benefits of this approach include: (i) avoiding publication bias (Perry 2022), with null results and preliminary findings more likely to be represented, and (ii) more accurately reflecting research of the time, as the time lag introduced by the publication process is avoided. These benefits are particularly important when trying to understand changes in research trends, the emergence of new methodologies, shifts in research priorities and identifying research gaps. As part of this synthesis, we characterize the salmonid research community, the biological systems studied, the research disciplines and methodologies employed, and the main questions that have been addressed to date, as well as emerging research trends. By doing so, this review offers a unique perspective on how salmonid research has evolved over the past two decades and gives insights into how emerging research trends and research gaps may shape the future of salmonid research.

## 2 METHODS

### 2.1 Data collection

Abstract books were downloaded from the NoWPaS website (https://nowpas.wordpress.com/previous-workshops/; n = 18) or acquired from previous attendees (n = 3) and covered the first NoWPaS meeting in 2005 up until the 2025 meeting (n = 21). Once downloaded, authors were randomly assigned for a series of years to extract information from the corresponding abstract book. An initial filter was conducted on the collated abstracts to remove those that were based on promoting a product or service as well as those presented by keynote speakers invited to the workshops (n = 78).

For the academic abstracts presented by master students, doctoral and postdoctoral researchers (n = 555), three authors per year independently extracted the following information: the research discipline(s) (Table S1), the method(s) used (Table S2), whether the study was tank or field based (or both), how theoretical or applied the topic was (scale from 1 to 5, respectively) as well as the main driver(s) of the research (*i.e*., *Fundamental research*, *Climate change*, *Management and restoration*, *Stocking*, *Hydropower and dams*, *Aquaculture*). For the theoretical-applied score, an average was taken across the three authors assigning a score. For the other categories, a fourth author assessed all assignments made by the three authors and made a final decision for each category.

Information for some categories that are not subjective to personal interpretation was extracted by one author. These included: country, or countries, of study (or if not available, the country where the author was based), study species, lifestage(s), habitat(s) and questions posed in the abstract. Gender was assigned using first names and the *genderizeR* package in R (Wais, 2016). An assessment of whether a NoWPaS delegate was still in academia in 2025 was also conducted based on an internet search. Affiliations were classified into six categories: academia, government, non-governmental organization (NGO) or non-profit, consulting and a more generic industry classification. Some attendees were not found in the internet search, or their online profile was not current, and were therefore classified as “unknown”. This classification was also used for attendees who had passed away.

### 2.2 Data analysis

In the following data analyses, when multiple responses or entries were indicated for a given variable in an abstract, each response or entry was treated as a separate occurrence.

#### 2.2.1 Research community

The gender balance of meetings was assessed using a gender ratio (number of males divided by number of females). Academic retention was assessed with a binomial generalised additive model (GAM) constructed using the *mgcv* package in R (Wood, 2011). Presence and absence from academia at the time of data collection was used as a binary response variable. The model contained a single smoothing term representing the number of years between the meeting date and the year of data collection (2025), referred to as “years since the meeting”, as well as an interaction term with gender. To avoid bias introduced by attendees of multiple meetings, if a full name appeared in more than one year, the earliest year of attendance was used. In addition to the full GAM, a second GAM was also created using a subset of well represented countries (Canada, Denmark, Finland, France, Iceland, Ireland, Norway, Sweden and UK) which contained the smoothing terms years since the meeting with an interaction term with gender and years since present day with an interaction term with country of affiliation.

#### 2.2.1 Co-occurence of categorical variables

Pairwise associations among categorical variables were explored using co-occurrence matrices and standardized Pearson residuals. To ensure sufficient representation across categories, analyses involving species were restricted to the six most frequently studied species and migratory forms (*i.e*., Atlantic salmon, landlocked Atlantic salmon, brown trout, sea trout, Arctic char and brook char), and trait analyses were restricted to categories with adequate occurrence frequencies (see section 3.7). For each variable pair, category co-occurrences were summarized in contingency tables and compared with frequencies expected under independence. Standardized Pearson residuals were then used as an exploratory measure of deviations from expected frequencies, with positive and negative values indicating over- and under-represented category combinations, respectively.

#### 2.2.2 Temporal patterns among research disciplines, methods and drivers

To assess temporal trends in the representation of research disciplines, methods, and drivers of studies, we did not use relative proportions or multinomial models because the different categories were not mutually exclusive (*i.e.*, a single abstract could be assigned to multiple categories). Although the total number of abstracts varied among years (mean = 26.4; SD = 6.6), it showed no significant linear trend over time (linear model, *lm*, using the *stats* package, version 4.6.1; *p* = 0.93). We therefore visualized temporal trends using annual counts of each category. For statistical analysis, we accounted for interannual variation in the number of abstracts by fitting generalized linear models with a quasi-binomial distribution. For each category representing at least 1% of all abstract assignments, we separately modelled the number of occurrences as a function of year (scaled to mean zero and unit variance), using the total number of abstracts each year as the number of trials. Because temporal trends were assessed independently for multiple categories, *p*-values were adjusted for multiple comparisons using the Benjamini-Hochberg (BH) correction. We also tested for differences in temporal trends between the two most studied species, *i.e.,* Atlantic salmon (*Salmo salar)* and brown trout (*Salmo trutta*, including the migratory form).

#### 2.2.3 Research positioning

To assess the scoring of research orientation along the theoretical (1) to applied (5) continuum, we first tested for potential reviewer bias using a linear mixed-effects model (*lmm*, package *lme4* in R, version 2.0-1). Since no significant reviewer effect was detected (the variance attributable to the three reviewers was negligible, < 0.01), we used the average score among the three reviewers to analyze research positioning (theoretical *vs.* applied) according to countries, disciplines, methods and species. Linear models (*lm*, package *stats* version 4.6.1) were employed on annual mean values to detect potential interannual tendencies. Note that we imposed a minimum threshold of n = 3 observations of any given variable per year. This criterion ensured sufficient occurrences to support robust inference on interannual patterns while limiting bias arising from sparsely represented categories. Finally, BH corrections were also applied to adjust *p*-values for multiple comparisons.

#### 2.2.4 Diversity of scientific questions

To analyze the evolution of questions posed in the abstracts, we conducted a temporal lexical analysis to quantify changes in vocabulary structure over time. Textual responses were tokenized into individual words using a standard bag-of-words approach, using *tidytext* R package (version 0.4.3). Common stop words and non-informative tokens (*e.g*., “what”, “the”, “is”) were removed to retain meaningful lexical units. For each year, word frequencies were computed and normalized to obtain relative proportions. These distributions were then used to estimate three complementary diversity metrics: Shannon diversity (H), lexical richness (S; number of unique words), and evenness (J). Temporal trends in these indices were assessed using linear models (*lm*, package *stats* version 4.6.1), and associated *p*-values were adjusted for multiple comparisons using the BH procedure. All statistical analyses were performed using R (version 4.6.1).

## 3 RESULTS

### 3.1 Research community

Gender ratio (male:female) has decreased over time, starting at around 2:1 in the early years of the conference (2005 – 2008; ignoring the anomalously high male representation in 2007) to the ratio tending towards 1:1 since 2015 (Figure 1a). The retention of researchers in academia was also assessed, which showed that the probability of attendees remaining in academia decreased as years since the meeting increased (χ2 = 16.79, *p* < 0.01) (Figure 1b). There was no significant interaction term between years since the meeting and gender, for either male (χ2 < 0.001, *p* = 0.5) or female (χ2 = 0.73, *p* = 0.42). Using a subset of the data which contained attendees with affiliations to well represented countries which contained at least 10 studies (Canada, Finland, France, Iceland, Ireland, Norway, Sweden and UK), there was no significant interaction between attendee’s country or gender influencing the relationship between their probability of remaining in academia and the number of years since the meeting.

**Figure 1:**
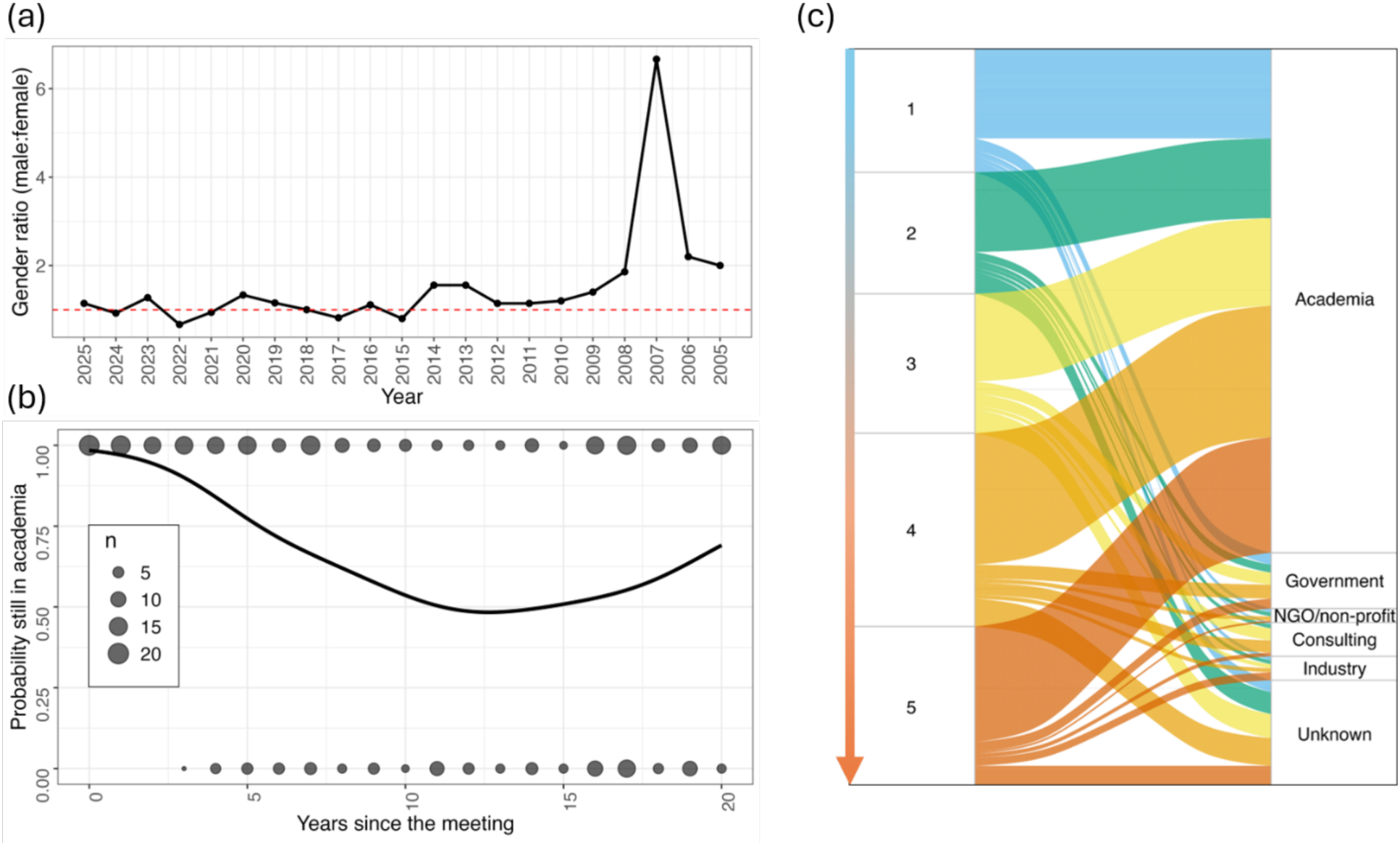
Plots showing how the NoWPaS community has changed over time. (a) Gender balance for all NoWPaS meetings, covering 2005 to 2025. (b) Probability of an attendee in a particular year remaining in academia at the time of data collection (2025). The black line shows estimates from a binomial generalized additive model (GAM) and the size of the circles show the raw frequency of those still in academia (1) and those that have left academia (0). (c) Sankey diagram which shows the affiliation of NoWPaS attendees at the time of data collection (right column), as well as the classification of their work presented at NoWPaS (1 = theoretical, 5 = applied).

All attendees of the three most recent conferences were still in academia, with the half-life of academics being 11 years (Figure 1b). Interestingly, after 13 years, the probability of attendees remaining in academia increased, peaking at 70% after 20 years. For those leaving academia, most moved to positions in government, or government adjacent agencies, as well as consulting and other industry positions; a process which was not heavily influenced by how applied the attendant’s research was (Figure 1c).

### 3.2 Geographical areas covered by the NoWPaS studies

Research was conducted in at least 26 countries since 2005 (Figure 2), but predominantly in Europe, notably in the United Kingdom (n = 128), Sweden (n = 76), Norway (n = 73), Finland (n = 61), France (n = 43), and Ireland (n = 31). In Europe, research was also conducted in Iceland (n = 16), Lithuania (n = 16), Denmark (n = 13), Spain (n = 8), Germany (n = 7), Greenland (n = 5), Switzerland (n = 4), Portugal (n = 4), Italy (n = 3), Estonia (n = 3), Belgium (n = 3), the Svalbard Archipelago (n= 1), and Poland (n = 1). In North America, research was conducted in Canada (n = 72) and the United States of America (n = 1) and focused primarily on the watersheds draining into the Atlantic Ocean. Some research was also conducted in the Southern Hemisphere, notably in the Kerguelen Islands (n =1), Falkland Islands (n = 1), Chile (n = 1), and Australia (n = 1). Overall, 71% of the research was conducted in the field, with just 29% conducted in “tank” based experiments (including *in silico*).

**Figure 2:**
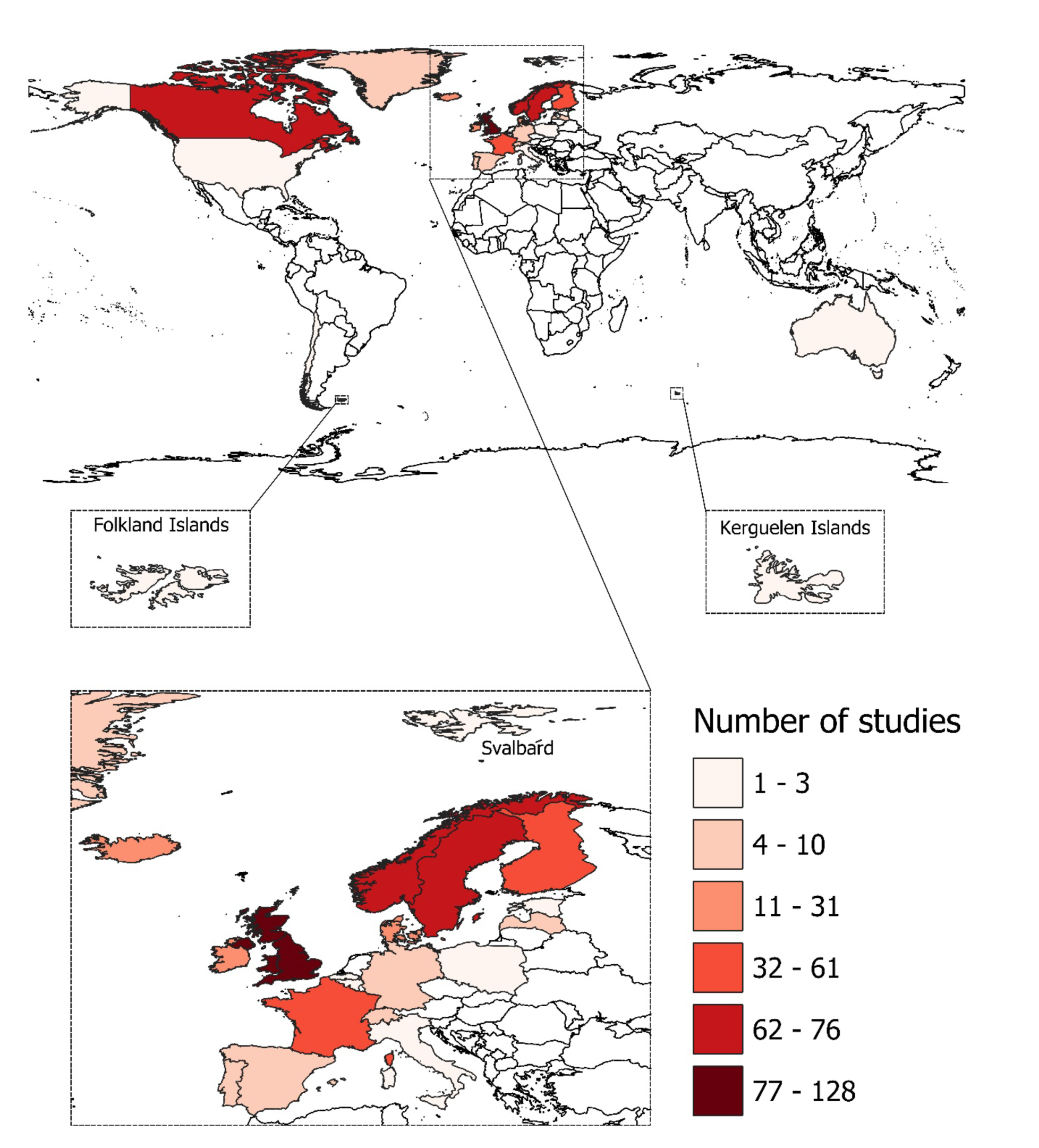
Countries in which NoWPaS research (excluding keynote speakers) was undertaken between 2005 and 2025.

### 3.3 Species (or groups), lifestages, and habitats

#### Species (or groups)

The abstracts submitted up to 2025 covered at least 5 salmonid genera, including *Salmo*, *Salvelinus*, *Oncorhynchus*, *Coregonus*, and *Thymallus* (Table S3). The genus *Salmo* was the best represented within the network and was investigated on 542 occasions. Within this genus, Atlantic salmon (*Salmo salar*) appeared 346 times, 17 of which focused on its landlocked form, and was studied consistently across the years (Table S3, Figure S1). Brown trout (*Salmo trutta*) was also consistently studied across years, yet most studies occurred before 2011, and was studied on 193 occasions, 57 of which targeted sea trout (Table S3, Figure S1). The marble trout (*Salmo marmoratus*) appeared on 3 occasions (Table S3, Figure S1).

The *Salvelinus* complex appeared in 50 abstracts (Table S3). Arctic char (*Salvelinus alpinus*) was the most represented with 29 studies, 11 of which focused on its anadromous form, yet appeared consistently only from 2018 onwards (Table S3, Figure S1). Brook char (*Salvelinus fontinalis*) was studied on 15 occasions, 2 of which focused on its anadromous form, and appeared sporadically across the years, with the anadromous form showing up for the first time in 2025 (Table S3, Figure S1). Dolly varden (*Salvelinus malma*) appeared 4 times between 2020 and 2023, while lake char (*Salvelinus umbla*) and lake trout (*Salvelinus namaychus*) only once both, with the latter being investigated as an invasive species in European freshwater lakes (Table S3, Figure S1).

The genus *Oncorhynchus* appeared sporadically before 2020 and became more consistent since then (Figure S1). In total, the genus was the focus of 13 abstracts, with rainbow trout (*Oncorhynchus mykiss*) appearing in 7 of them, coho salmon (*Oncorhynchus kisutch*) appearing 4 times, and chinook (*Oncorhynchus tshawytscha*) and pink (*Oncorhynchus gorbuscha*) salmon appearing both once (Table S3). In 2025, pink salmon was investigated as an invasive species outcompeting Arctic char and Atlantic salmon in the North Atlantic Ocean. The genera *Coregonus* and *Thymallus* were the focus of 6 abstracts, with the European grayling (*Thymallus thymallus*) occurring 3 times, the Atlantic (*Coregonus huntsmani*) and European (*Coregonus lavaretus*) whitefish both once in 2025, and on one occasion the whitefish species was not specified.

On 63 occasions and consistently across the years, the NoWPaS network studied salmonids in conjunction with other organisms (Table S3, Figure S1). These studies evaluated gut microbiome, parasites (*e.g.*, pearl mussel, sea lice) but also prey-predator relationships with both freshwater (*e.g*., burbot, pike) and saltwater (*e.g*., striped bass) fish, and the interactions with aquatic vegetation, algae, macrozoobenthos, and mammals such as beavers and otters.

#### Lifestages

The NoWPaS research up to 2025 covered the entire lifestage continuum of salmonids, including egg, alevin, fry, juvenile, smolt, post-smolt, adult, carcass and kelt stages (Figure S2). We note that the juvenile stage (as reported in several abstracts) can also include other potential early lifestages such as fry or parr. Except for the alevin, post-smolt, kelt and carcass stages, all other lifestages were investigated consistently across years (Figure S2), with the adult stage being the most represented in the NoWPaS research (n = 157), and the carcass stage only investigated once in 2025.

Our results suggested that salmonid species received uneven attention across lifestages (Figure S3). For instance, Atlantic salmon was mostly associated with smolt and post-smolt stages but underrepresented before smoltification, whilst landlocked Atlantic salmon showed a clear dichotomy between the well-represented kelt stage and the underrepresented juvenile stage. Brown trout was mostly associated with early lifestages, whereas research on sea trout mostly focused on smolts and kelts rather than other lifestages.

#### Habitats

The abstracts submitted up to 2025 focused on at least 9 habitats/environments (Figure S4). The riverine environment was the most represented within the network, with rivers consistently studied across years (n = 377), whereas frozen rivers were only studied five times (up to 2014) and the entire watershed just once. The lacustrine environment was also consistently investigated within the network but only accounted for 34 abstracts. The coastal, estuary, and fjord environments were studied 4, 6, and 7 times, respectively, and their presence in abstracts was sporadic across years, though the fjord environment became regularly present from 2021. The marine environment was regularly investigated within the network and accounted for 68 abstracts. Finally, controlled environments, notably farms and hatcheries, were present sporadically. Similarly to species-lifestage combinations, our results showed that habitats were unevenly represented across salmonid species (Figure S3). While, overall, the number of studies in marine environments was much lower than in freshwater environments (Figure S2), of those studies conducted in marine environments, Atlantic salmon and sea trout were well-represented. Consistent with their life-history strategies, research on Arctic char and landlocked Atlantic salmon was mostly associated with lacustrine environments and underrepresented in rivers (Figure S3).

### 3.4 Research disciplines of NoWPaS studies

Across all years combined, disciplines were dominated by *Ecology*, which accounted for 23% of all abstract assignments (Figure 3a). *Genetics and genomics*, *Movement ecology*, and *Physiology* each represented 13% of assignments, although they were associated with different focal species (Figure S3). *Genetics and genomics* were positively associated with Atlantic salmon and Arctic char, *Physiology* was positively associated with Atlantic salmon only, and *Movement ecology* was strongly well-represented in studies of sea trout (Figure S3). This was followed by *Hydro-geomorphology* (10%), which was associated with brown trout and brook char, and *Evolutionary biology* (9%), which was associated with Arctic char. *Habitat use* and *Behavioural ecology* each accounted for 7% of abstract assignments, whereas *Disease and parasitology* (4%) and *Ecotoxicology* (1%) were comparatively rare.

**Figure 3:**
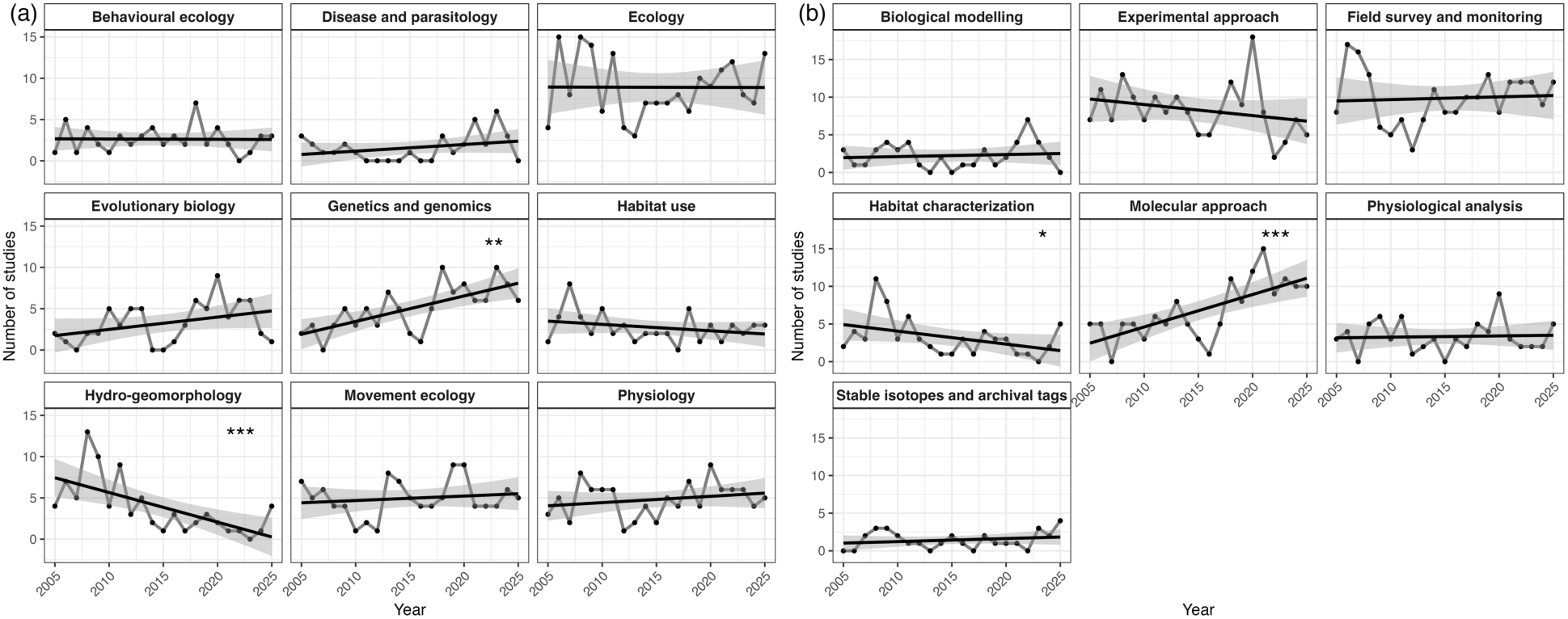
Temporal change in the occurrence of a) research disciplines and b) methods. Points represent the annual number of studies per category. Black solid lines show linear regression fits for each category, with shaded areas indicating 95% confidence intervals around the fitted relationships. Asterisks denote statistically significant temporal trends based on the slope of the linear regression (*p* < 0.05*, *p* < 0.01**, *p* < 0.001***).

Temporal trends differed among those disciplines (Figure 3a, Table S4). Notably, there was a significant increase in the representation of the *Genetics and genomics* discipline over time (*p* < 0.01), although this trend was driven by studies on Atlantic salmon while studies on brown trout (including sea trout) did not show this pattern (Figure S5; Table S5). In contrast, the *Hydro-geomorphology* discipline showed a significant decline through time (*p* < 0.001) which was consistent across both species (Figure S5; Table S5). There was no change in the overall representation of the *Ecology* discipline, despite a significant decrease among studies on brown trout (Figure S5; Table S5). No significant temporal trends were detected for all the other disciplines, indicating that their representation remained broadly stable over the study period despite interannual variations.

### 3.5 Methodological changes through time

Overall, *Field survey and monitoring* methodologies (28%) and *Experimental approaches* (23.5%) were more frequently implemented, which were positively associated with studies of sea trout and brown trout, respectively (Figure S3). This was followed by *Molecular approaches* (19%), which were well-represented in studies of Arctic char and Atlantic salmon. They also significantly increased in occurrence over time (p < 0.001; Figure 3b; Table S6), although this trend was driven by studies on Atlantic salmon while studies on brown trout (including sea trout) did not show this pattern (Figure S5; Table S7). Less commonly used methods included *Physiological analysis* (9%), *Habitat characterization* (9%), *Biological modelling* (6.5%), *Stable isotopes and archival tags* (4%), and *Literature review* (1%). *Habitat characterization* methods were positively associated with studies of brown and brook char and showed a significant decrease in occurrence over time (*p* < 0.05) which was consistent across both main species (*i.e.,* Atlantic salmon and brown trout; Figure S5; Table S7). No significant temporal trends were detected for the other methodologies (Figure 3b; Table S6), except for a significant decrease in *Experimental approaches* on brown trout (Figure S5; Table S7).

Within broad methodological categories, the use of more specific methodologies also changed over time, particularly with the emergence of new technologies. For example, *molecular* methodologies evolved substantially (Figure 4), with early studies dominated by classical genetic markers such as allozymes and microsatellites, before progressively shifting towards SNPs-based analyses. Gene expression studies also showed a methodological transition, moving from targeted gene expression methods to RNA sequencing. More recent years were characterized by a strong diversification of genomic methods, with the emergence of GWAS, eDNA, CRISPR gene editing, metabarcoding, epigenomics, genomic offset, reflecting a transition toward the omics era. Within *Field survey and monitoring* approaches (Figure S6a), morphological measurements (*e.g.*, growth-related traits) remained consistently and widely used throughout the last 20 years, alongside electrofishing. Telemetry methods also evolved over time, with a progressive shift from radio telemetry toward acoustic telemetry, while PIT tags were used consistently over time. For *Physiological analysis*, the use of respirometry remained fairly constantly applied through time (Figure S6b). Starting in 2019 there were also several methods only applied once, perhaps demonstrating novel emerging methodologies. In addition, starting in 2016, there was a burst of gamete analyses. *Habitat characterization* was dominated by broad habitat modelling, assessment, and classification techniques, with hydraulic modelling prevalent in the first decade and remote sensing and hydraulic measurements in the second (Figure S6c). Among *Experimental approaches*, there was a slight transition from field and channel experiments to lab experiments (Figure S6d). *Biological modelling* methods display new methodologies with the appearance in the later years of the use of machine learning and Dynamic Energy Budget Modelling (Figure S6e). Finally, regarding *Stable isotopes and archival tags*, a recent use of biogeochemical tools (*e.g.*, scale, otolith and eye lenses chemistry) could be detected within abstracts, while stable isotopes have been consistently used over the past 20 years (Figure S6f).

**Figure 4:**
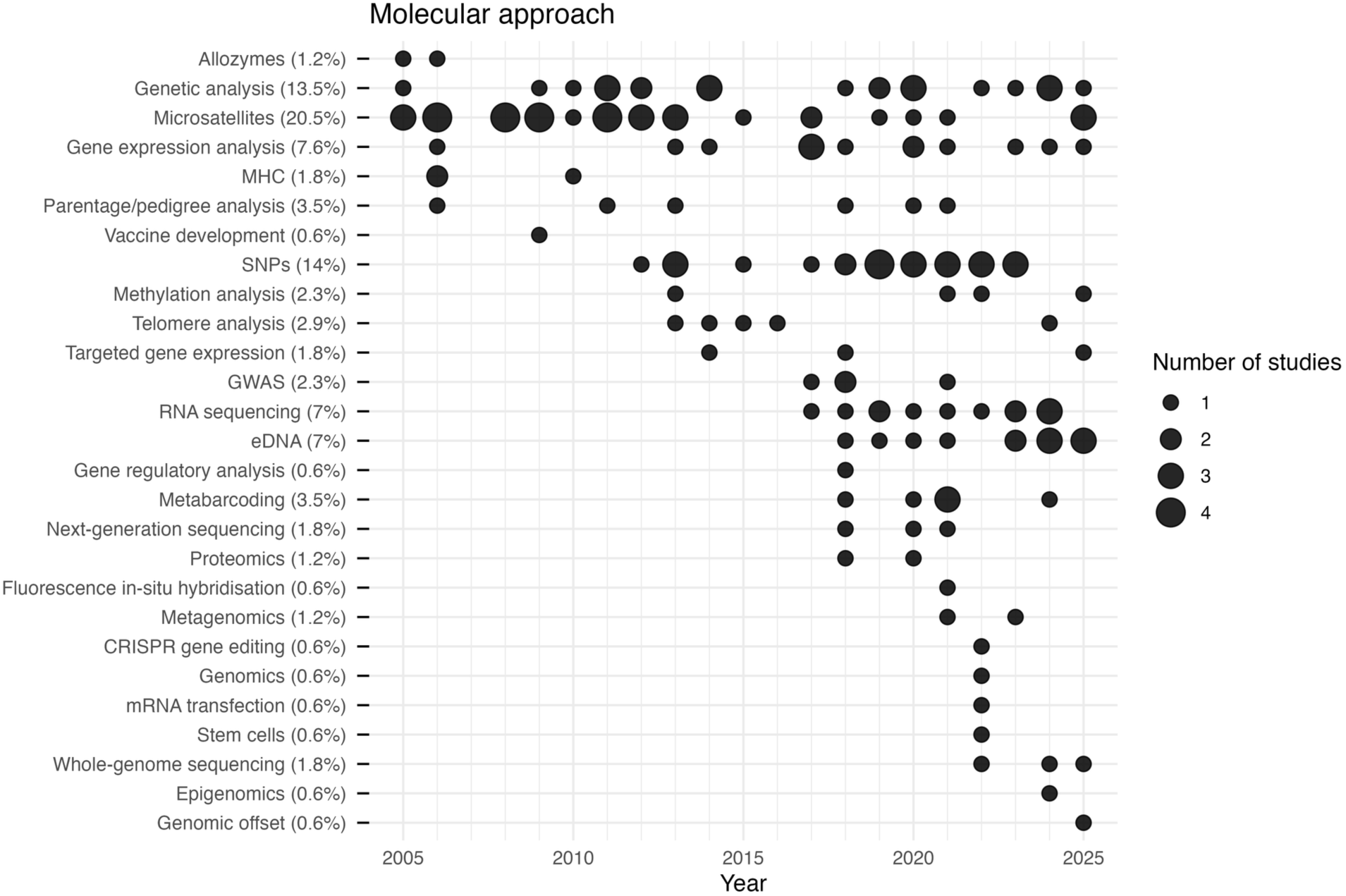
Temporal diversification of methods within *Molecular approaches*.

### 3.6 Research positioning of studies and associated drivers

Over the 2005–2025 period, research was, on average, slightly more applied than theoretical (mean = 3.16, SD = 1.34; 3 being the median between theoretical and applied), with no clear temporal trend. However, variation in positioning along the theoretical–applied continuum was observed across countries, disciplines, methods, and study species. At the country level (Figure 5), relatively stable temporal patterns were found in Canada (3.54 ± 1.43), Denmark (2.95 ± 1.59), Finland (3.06 ± 1.61), France (2.45 ± 1.43), Ireland (3.61 ± 1.36) and Sweden (3.47 ± 1.61), whereas an increasing shift toward more applied research was observed in Iceland (3.25 ± 1.59, *p* = 0.009), Norway (3.02 ± 1.52, *p* = 0.054) and the United Kingdom (3.04 ± 1.55, *p* = 0.007). Similarly, disciplinary patterns revealed stable orientations toward theoretical research in *Evolutionary biology* and *Physiology*, and toward more applied research in *Hydro-geomorphology* and *Movement ecology* (Figure 5). In contrast, *Behavioural ecology* (not significant) and *Ecology* (*p* = 0.03) showed a gradual shift toward more applied research over time. Comparable increasing trends were also evident for certain methodological approaches, particularly studies using stable isotope techniques and physiological analysis (Figure S7). Finally, species-specific patterns indicated that research on Atlantic salmon and brown trout tended to occupy intermediate positions along the continuum, whereas studies focusing on brook char, landlocked Atlantic salmon, and sea trout were generally associated with more applied research orientations (Figure S8).

**Figure 5:**
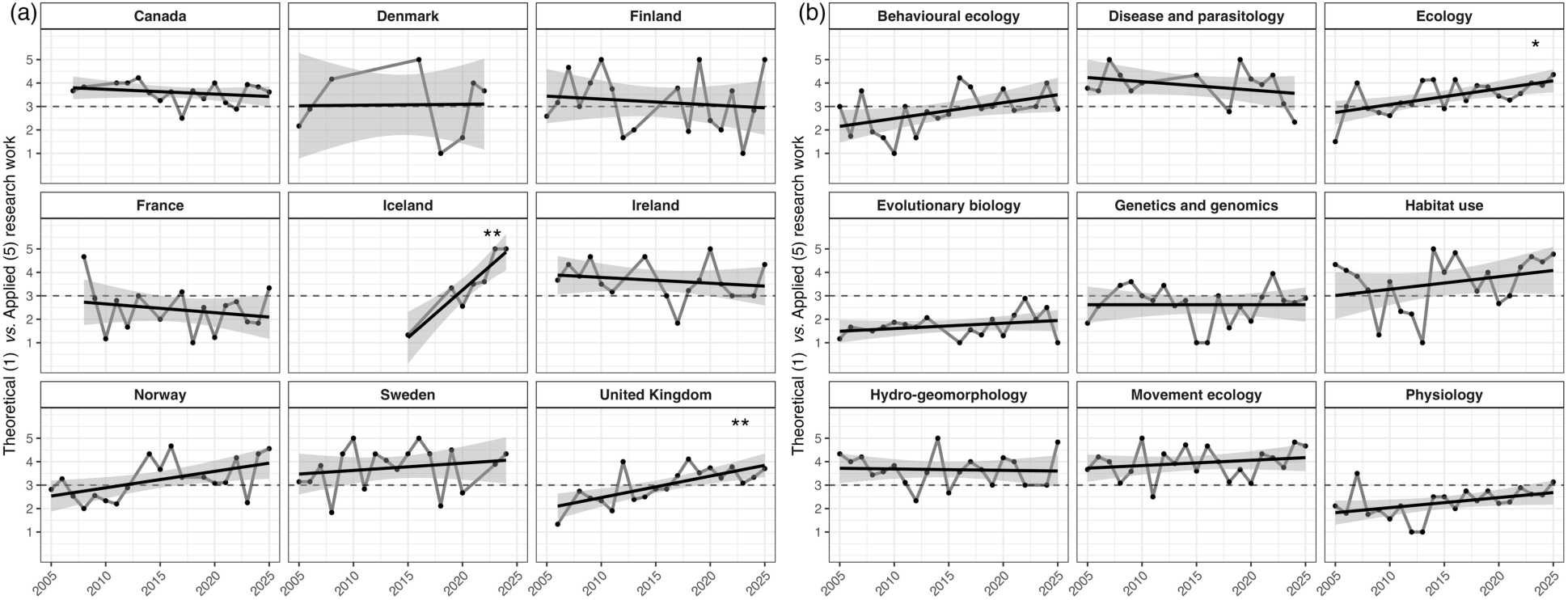
Temporal change of research positioning scores (theoretical: 1; applied: 5) according to a) the country of study and b) research disciplines. Points represent the annual average score obtained for each category. Black solid lines show linear regression fits for each category, with shaded areas indicating 95% confidence intervals around the fitted relationships. The dotted line represents a neutral value. Asterisks denote statistically significant temporal trends based on the slope of the linear regression (*p* < 0.05*, *p* < 0.01**, *p* < 0.001***).

Overall, research was most frequently motivated by *Fundamental research*-oriented objectives, which accounted for 42% of all assignments, followed by *Management and restoration*-related motivations (28%; Figure 6a). The latter were well-represented in studies of resident brown trout and brook char (Figure 6b). *Climate change* (10%) was positively associated with studies of Arctic char, whereas *Aquaculture* (7%) was well-represented in studies of Atlantic salmon. More applied study drivers, namely *Hydropower and dams* (9%) and *Stocking* (4%), were well-represented in studies of the landlocked form of Atlantic salmon, as well as sea trout (*Hydropower and dams* only). The overall representation of *Fundamental research* and *Management and restoration* motivations did not change significantly over time (Figure 6a; Table S8), although there was a significant decrease in *Fundamental research*-oriented studies on brown trout (including sea trout) (Figure S9; Table S9). In contrast, *Climate change* (10%; *p* < 0.01) and *Aquaculture* (7%; *p* < 0.05) were study drivers that increased significantly in occurrence over the study period, in both main species and in Atlantic salmon only, respectively (Figure S9; Table S9). Finally, *Hydropower and dams* (9%), and *Stocking* (4%) did not exhibit significant temporal trends.

**Figure 6:**
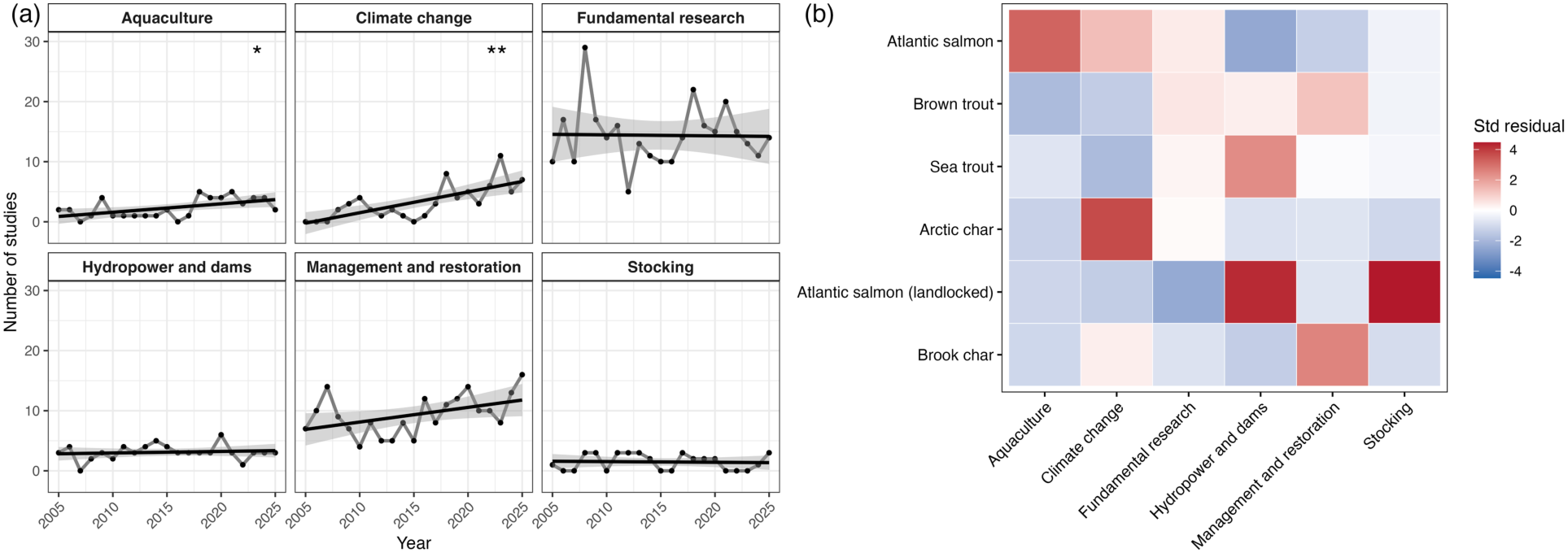
a) Temporal change in the occurrence of study drivers. Points represent the annual number of studies per category. Black solid lines show linear regression fits for each category, with shaded areas indicating 95% confidence intervals around the fitted relationships. Asterisks denote statistically significant temporal trends based on the slope of the linear regression (*p* < 0.05*, *p* < 0.01**, *p* < 0.001***). b) Co-occurrence of study drivers and study species measured by standardized Pearson residuals from chi-squared tests of independence. Positive residuals (red) indicate more frequent combinations than expected under independence, while negative residuals (blue) indicate under-represented combinations during the 2005-2025 period. Std residuals: Standardized Pearson residuals.

### 3.7 Common traits measured in salmonid studies

A wide range of traits were measured across studies, although several were repeatedly assessed (Figure 7; Table S10). *Growth* was the most frequently reported trait, appearing in 12% of assignments, followed by *Movement* and *Survival*, each mentioned in 9% of the assignments. Other individual-level traits were less commonly measured, including *Behaviour* (6%), *Metabolism* (3%), *Diet* (3%), *Morphology* (2%), *Timing* of life-history events (2%; *e.g*., migration timing), *Stress* (1%), *Infection rate* (1%), *Age at maturity* (1%), and *Straying* (1%). Genetic-related traits included *Genetic structure or diversity* (6%), *Gene expression* (3%), as well as *Introgression* (1%). Habitat-related variables were also commonly reported, with *Habitat use* and *Habitat characteristics/quality* each occurring in 5% of assignments. At the population level, *Density and abundance* were reported in 5% of assignments. All remaining traits were reported in fewer than 1% of the assignments (Table S10).

**Figure 7:**
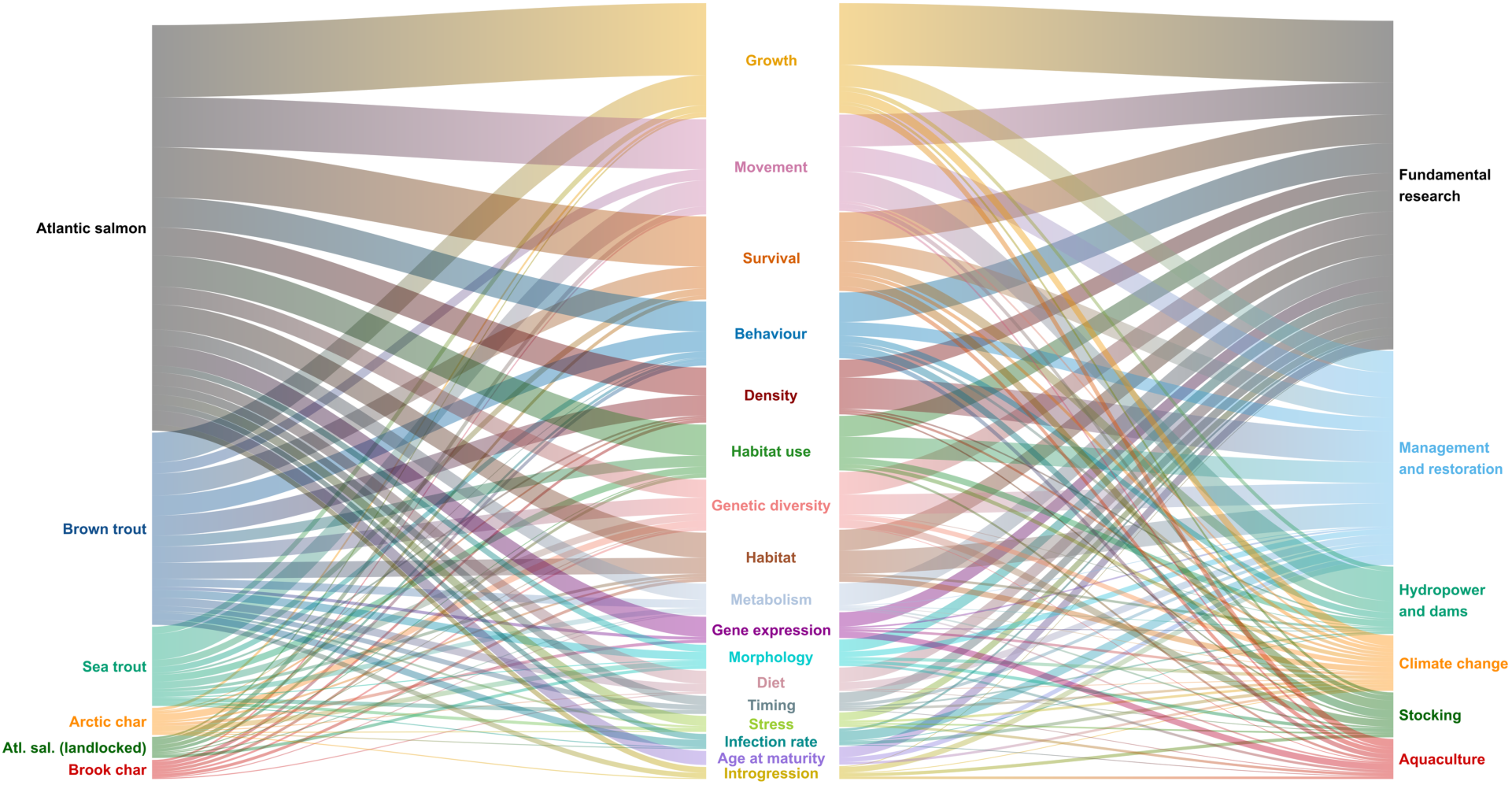
Sankey diagram illustrating the co-occurrence of study species (left), measured traits (middle), and study drivers (right). Line thickness is proportional to the number of combinations. Only the six main species and traits reported in more than 10 occurrences are shown. Atl. sal. (landlocked): Atlantic salmon (landlocked).

Several traits showed clear associations with particular species and study drivers (Figure 7). *Age at maturity*, for instance, was reported exclusively in studies on Atlantic salmon, which was also more frequently associated with *Gene expression* measures and with *Aquaculture*-related research. *Behaviour*al traits, by contrast, were well-represented in studies on brook char and brown trout, while *Density* and *Habitat characteristics/quality* were more commonly associated with these same two species and with *Management and restoration* motivations. *Morpholog*ical traits were strongly tied to studies on Arctic char, whereas *Movement*-related traits were strongly associated with sea trout and *Hydropower and dams* drivers. Finally, *Introgression* was positively associated with *Stocking*-related drivers, and *Metabolism* traits were linked to *Fundamental research*.

### 3.8 Evolution of scientific questions

Lexical diversity (H) and richness (S) in research questions increased over the study period, whereas lexical evenness remained relatively stable (Table S11). Linear models indicated a significant positive temporal trend in Shannon lexical diversity (H; *p* = 0.04) and lexical richness (S; *p* = 0.05). In contrast, lexical evenness (J) did not vary significantly over time (*p* = 0.60). Interestingly, temporal patterns of H differed among research disciplines (Figure 8, but see Table S11 for temporal patterns of S and J by research discipline). Significant increases in Shannon diversity were observed in *Disease and parasitology* (*p* < 0.01) and *Genetics and genomics* (*p* < 0.001), with marginally non-significant increases in *Evolutionary biology* (*p* = 0.06) and *Movement ecology* (*p* = 0.09). Lexical richness showed similar patterns, increasing significantly in *Disease and parasitology* (*p* < 0.01) and *Genetics and genomics* (*p* < 0.01), while weaker positive trends were detected in *Evolutionary biology* (*p* = 0.06) and *Movement ecology* (*p* = 0.09). In contrast, *Hydro-geomorphology* exhibited negative, though non-significant, trends in both diversity (*p* = 0.10) and richness (*p* = 0.07). No significant temporal changes in lexical evenness were detected across disciplines after multiple-testing correction.

**Figure 8:**
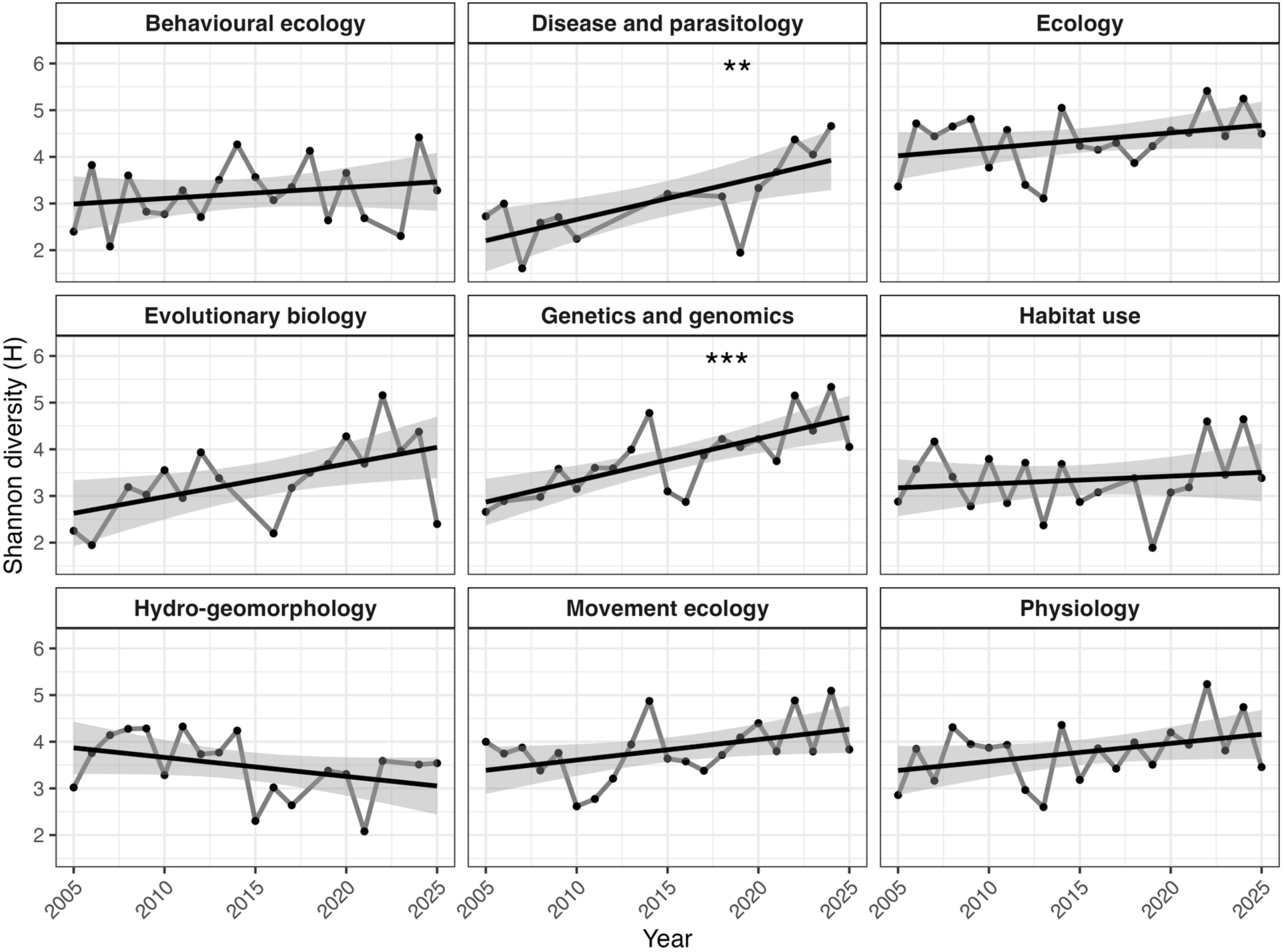
Temporal variation in Shannon lexical diversity (*H*) by research discipline calculated from tokenized abstract research questions. Points represent the annual value of the index. Black solid lines show linear regression fits for each index, with shaded areas indicating 95% confidence intervals around the fitted relationships. Asterisks denote statistically significant temporal trends based on the slope of the linear regression (*p* < 0.05*, *p* < 0.01**, *p* < 0.001***).

## 4 DISCUSSION

We used abstracts presented at NoWPaS workshops to synthesize research trajectories on salmonids over the past two decades. This approach provides a complementary research perspective to peer-reviewed publications alone, as it captures preliminary findings, emerging ideas, and non-published results. We characterized changes in research themes, study systems, methodological approaches, and disciplinary contributions within the research community. The following discussion addresses these main dimensions, situating them within the broader context of current salmonid research, before considering their implications in terms of knowledge gaps and future research directions.

### 4.1 Insights through the NoWPaS lens

The synthesis presented here reflects the scope and composition of the NoWPaS community and should therefore be interpreted through this lens. It primarily captures research conducted and presented by ECRs and is geographically biased toward the Northern Hemisphere and the Atlantic, where all workshops have been held. The location of NoWPaS research is skewed towards the countries where the workshop has been held more frequently (*e.g.*, the United Kingdom; Figure 2). There may also be a positive feedback loop as ECRs transition into established researchers who promote the network among the ECRs they (co-)supervise. Consequently, NoWPaS research and its geographic distribution only reflect a fraction of global salmonid research. Research from Asia, Western North America, and South America is almost completely underrepresented. However, by providing a structured and quantitative overview of research presented within this international ECR network, this review highlights key trends, gaps, and emerging themes which are likely to be relevant across all salmonids.

### 4.2 Past and present trends in salmonid research

#### 4.2.1 Composition of the community

Gender parity of attendees has improved since the early years of the conference, especially after 2010, reflecting the trend seen at other larger fish focused symposia (Kaufmann and Perry, 2026) or in other scientific conferences and meetings (Piccoli and Guidobaldi, 2021). This trend shows the success of strategies implemented by organizing committees and scientific societies to improve gender parity (*e.g*., Penaluna et al., 2017). The trend is also consistent with general observations on gender inequality within and beyond fisheries sciences (Wilkes et al., 2016; Penaluna and Arismendi, 2022; Basilio Hazas et al., 2026). In contrast to major scientific societies (*e.g*., FSBI, AFS), NoWPaS is substantially less known within the fisheries community, and ECRs typically become aware of it through word of mouth, often from their direct supervisors or past attendees. As a result, recruitment relies on a relatively small and interconnected participant pool rather than broad community-wide visibility. Consequently, any existing demographic imbalances within that pool are more likely to persist, potentially reinforcing the underrepresentation within the network. In addition, the methodology used here to assign gender based on first name has its limitations, however, as this data is not available, it provides an imperfect working proxy for assessing balance in binary genders through time.

Our analysis did not suggest significant influence of gender on the probability of remaining in academia after having attended a NoWPaS meeting. As expected, attendees of the most recent meetings (2023-2025) were almost exclusively still in academia, reflecting their early-career status. Among attendees of meetings held between 2011 and 2022, the probability of remaining in academia declined to ∼ 50%. Interestingly, this probability was higher for the earliest conference cohorts (2005-2010), reaching ∼ 70% for attendees of the 2005 meeting. Although our data does not allow us to determine the cause of this pattern, it may reflect changes in the academic job market over the past two decades. The availability of permanent academic science, technology, engineering, and mathematics (STEM) positions declined or remained stable relative to the growing number of PhD students, leading (or forcing) an increasing proportion of ECRs to pursue careers in government, NGOs, consulting, or industry (Hansen et al., 2014; Hampton and Labou, 2017; Fetcher et al., 2019; Cheng, 2023). Interestingly, our results suggest that researchers are no more likely to leave academia based on how applied their research is.

#### 4.2.2 Changes in biological study systems through time

Our analysis showed that NoWPaS research was largely conducted in European countries and the Atlantic provinces of Canada. As such, the genus *Salmo* dominated NoWPaS research, with Atlantic salmon (including landlocked salmon) being the species most frequently studied, followed by brown trout (including sea trout). We speculate that this could be related to research priorities and funding within the European research landscape, ultimately reflecting society’s perception regarding Atlantic salmon (Kochalski et al., 2022). Our results showed that *Aquaculture* was a key driver of Atlantic salmon research, with research also focusing on molecular methods and traits such as gene expression and age at maturity, all of which increased in prevalence over time. At the same time, wild Atlantic salmon populations have experienced severe declines across much of their range, a trend increasingly compounded by *Climate change* and shifting marine ecosystem conditions, which also motivates a substantial body of research (Dugdale et al., 2016; Rinaldo et al., 2024). Beyond taxonomic imbalance, our results also revealed a persistent mismatch between life stages and habitats across species. Research on anadromous forms primarily addressed smoltification and adult stages present in rivers, where fish are easier to observe and sample, with marine environments understudied. Research on non-anadromous forms focused on earlier life stages, riverine and lacustrine environments (Hatanpää et al., 2025). While most research was focused on single species, a consistent body of work took an ecosystem-level approach, examining interactions between salmonid species and other taxa, including, more recently, mammals (Sortland et al., 2023; Macarthur et al., 2026).

#### 4.2.3 Conceptual and methodological changes in research through time

Our results revealed a parallel increase in the prevalence of the *Genetics and genomics* discipline and *Molecular approa*ches over time, whereas the *Hydro-Geomorphology* discipline and methods of habitat characterization showed a corresponding decline over the same period. These shifts likely mirror a broader change in research priorities, paralleling the growing prominence of *Aquaculture* and *Climate change* as drivers of research within the network.

The reorientation toward molecular and genomic tools is likely driven by technological innovations. Allozymes were present in two of the earliest meetings, but not seen again, showing the broader decline of this methodology. In the period 2005 to 2011, microsatellites were in their prime and were the main molecular method of choice. However, from 2012 this begins to change, and a radiation of molecular methods start to appear due to the widening accessibility of high-throughput sequencing to researchers, due to factors like decreasing costs; the use of single nucleotide polymorphisms (SNPs) being a good example of this. A similar pattern in the global use of genetic markers has been documented elsewhere (Carvalho, 2025; Sturrock et al., 2026b). After 2017, high-throughput sequencing methodologies were being applied to a wide variety of applications such as transcriptomics and RNA sequencing (Wang et al., 2009), suggesting that research is developing from understanding what genetic diversity is present to how that diversity is expressing itself (beyond single genes). In addition to this, an increase in environmental DNA was also observed (Yates et al., 2021; Perry et al., 2025; Macarthur et al., 2026), an approach which has harnessed the power of molecular approaches. The increase is not surprising given that it is non-invasive, cost- and time-efficient when applied at scale, and has the potential to detect fish at low densities.

Another methodological shift is seen in the *Field Survey and Monitoring* discipline with the development in animal tracking. Specifically, Passive Integrated Transponder (PIT) tagging and radio telemetry have a consistent application over the time series, whereas acoustic telemetry has seen larger increases in the last 10 years. Given that acoustic telemetry can track fish across larger areas and provide richer movement data, it is understandable why it is becoming the method of choice for tracking the movement of salmonids. Not only this, but technological development into the miniaturisation of acoustic tags (Lennox et al., 2025), increasing battery length (Jacoby and Piper, 2025) and sharing acoustic tracking arrays (Lennox et al., 2024) mean that this approach can now be applied to a broader array of research questions.

More broadly, stable isotope analyses have remained a cornerstone of fish ecology over the past two decades because they provide robust information on trophic ecology, resource use, and habitat occupancy across multiple temporal scales. Recently, studies have increasingly combined stable isotope analyses with archival-tag approaches to integrate trophic and environmental histories, an approach that has proven effective in both non-salmonid (Teichert et al., 2022; Li et al., 2025) and salmonid fishes (Betts et al., 2026b).

Beyond research disciplines and methodologies, clear patterns also emerged in the traits measured across NoWPaS studies. Individual growth was the most frequently reported trait, likely reflecting both the relative ease of its measurement and its established value as an indicator of individual and ecosystem health (Gallagher et al., 2022). This was closely followed by survival and movement, the latter being tightly linked to sea trout studies, the rise of new telemetry methods and to large concern over hydropower and dams impeding connectivity of migrating species (Chan et al., 2025). Genetic-related traits, including genetic structure, diversity and gene expression, were also well represented, consistent with the growing prominence of molecular approaches described above. Behaviour also remained among the most frequently measured traits, reflecting its broad relevance across species and research contexts (Candolin and Wong, 2012). Density and habitat quality were commonly measured in the context of management and restoration of Brook char and Brown trout populations (*e.g*., Alcaraz-Hernández et al., 2016). Notably, morphological traits were tightly linked to studies on Arctic char (*e.g*., Jónsdóttir et al., 2024) and age at maturity was exclusively reported in Atlantic salmon, especially with recent studies focusing on the discovery of a large effect gene influencing maturation timing (Czorlich et al., 2022; Raunsgard et al., 2024; Maamela et al., 2025).

#### 4.2.4 Research positioning

Overall research orientation across the 21 NoWPaS conferences was intermediate (3.16 ± 0.63), indicating a balanced distribution between fundamental and applied studies in salmonid research. This is consistent with the dual role of salmonids as model organisms for addressing core ecological and evolutionary questions—such as the importance of life-history diversity and the role of metapopulation dynamics under environmental change (Fontaine et al., 2025; Lamarins et al., 2025; Sturrock et al., 2026a)—and as taxa of high conservation concern across much of their native range, which has motivated extensive restoration and rehabilitation efforts (Louhi et al., 2016; Bernthal et al., 2022; Koski et al., 2025; Fauchet et al., 2026; Moccetti et al., 2026). However, marked heterogeneity emerges when examining temporal trends across countries and disciplines.

At the country level, Iceland and the United Kingdom showed significant shifts toward more applied research, with a similar trend in Norway. This asymmetry suggests a broader reorientation of salmonid research priorities toward solution-driven approaches, likely linked to increasing environmental pressures and the consequent demand for actionable scientific outputs aimed at conservation and management. For example, Atlantic salmon populations in Norway are increasingly affected by recent biological invasions of pink salmon (*e.g*., Sandlund et al., 2019; Thorstad et al., 2025) and hybridization with domesticated fish (*e.g*., Perry et al., 2019; Diserud et al., 2022), while studies from the United Kingdom have emphasized the critical importance of improving smolt and post-smolt migration success to sustain population viability (Rodger et al., 2025; Adams et al., 2026). There has also been a concerted effort in breaking down silos in countries like the UK, with the FSBI’s 2026 symposium titled: ‘Breaking Siloes in Fish Biology’. One example of siloing is between institutions, like universities and government agencies/NGOs (who undertake more applied research), the erosion of which is likely to promote more applied research presented in academic conferences, like NoWPaS. Lastly, the change in the UK could also be due to a shifting funding landscape, with universities potentially supplementing stretched research budgets with income streams from non-government stakeholders who require more applied outputs; or due to shifts in UK Research and Innovation (a non-departmental public body of UK Government) increasing emphasis on application of research and less on curiosity driven research.

In opposition, France stands out as systematically more oriented toward fundamental research (2.45 ± 1.43, over 2005-2025), reflecting possible differences in research structuring and institutional roles. In France, applied salmonid research is often conducted within national or regional organizations (*e.g.*, Office Français de la Biodiversité, migratory fish management associations), whereas academic institutions tend to focus on mechanistic and theoretical questions, perhaps reflecting a more siloed approach. It may also reflect a participation bias, as French representatives attending these conferences were predominantly affiliated with institutions focusing on fundamental research topics, or the sparce viable populations of heavily managed salmonids like Atlantic salmon in France.

Research disciplines differed markedly in their average application scores. For instance, disciplines such as *Movement Ecology*, *Disease and Parasitology*, *Habitat Use*, and *Hydro-Geomorphology*, which showed more applied scores (Figure 5), are more directly connected to conservation and management challenges. Their outputs frequently inform operational decisions regarding fish passage and migratory connectivity (Calles and Greenberg, 2005; Östergren and Rivinoja, 2008; Forty et al., 2016; Goerig et al., 2016; Nyqvist et al., 2017), disease risk and survival (Werkman et al., 2011; Eldøy et al., 2020; Philpott et al., 2025), identification, understanding, and protection of spawning and nursery habitats (Palm et al., 2007; Louhi et al., 2008; Roncoroni and Lane, 2019; Koski et al., 2025; Roncoroni et al., 2026), or the preservation of thermal refugia under climate change (Dugdale et al., 2015; O’Sullivan et al., 2019, 2021). Notably, *Ecology* was the only discipline showing a significant increase in application scores over the 2005–2025 period. This likely reflects a growing emphasis on translating advances in ecological understanding into effective conservation and management strategies. As many salmonid species are currently classified as Vulnerable (*e.g*., Siberian taimen *Hucho taimen*, bull trout *Salvelinus confluentus*) or Endangered (*e.g*., Adriatic grayling *Thymallus aeliani*) by the International Union for Conservation of Nature Red List (IUCN), the scientific literature increasingly advocates for a closer integration of fundamental and applied research.

#### 4.2.5 Diversity of research questions

Lexical diversity and richness increased significantly over time, indicating a progressive expansion of vocabulary used in research questions (Figure 8, Table S11). However, evenness remained stable, suggesting that this increase is driven by vocabulary expansion rather than changes in word distribution structure. This was especially driven by two research disciplines: *Genetics and genomics* and *Disease and parasitology*. We hypothesize that this growing diversity of research questions reflects three complementary processes: (i) the geographical expansion of the NoWPaS community, (ii) the rapid development of new methods and analytical tools and (iii) scientific questions often shifting from describing patterns to explaining mechanisms and interactions as knowledge on a system accumulates. Over the last two decades, the symposium has progressively expanded from its Scandinavian origins toward Western Europe and North America, fostering interactions among researchers working in different ecological, evolutionary, and management contexts. This expansion likely promoted the inclusion of new research perspectives, study systems, and focal species. For example, species such as Arctic char and brook char, which hold particular ecological and cultural importance in North America, became increasingly represented in conference discussions. During the early years of NoWPaS (2005–2008), many studies focused on relatively specific and localized questions, *e.g.*, what are the effects of ice cover on habitat use and survival? In contrast, recent conferences have increasingly addressed broader and more integrative questions, *e.g*., how does variation in life-history strategies across populations and years influence the resilience and management of salmon under climate change? This shift coincides with the emergence and widespread adoption of powerful new approaches, including high-throughput sequencing, advanced tagging technologies, and increasingly sophisticated laboratory and analytical techniques.

Undoubtedly, methodological developments have expanded the scope of investigation from fine-scale biological and population-level processes to large-scale ecological and evolutionary questions. As illustrated by the growing prominence of genetics-related research (Figures 3 and 4), with tools initially used to characterize population structure and genetic diversity now being applied to address mechanistic questions related to adaptation, biodiversity monitoring, and ecosystem functioning. More generally, advances in data collection and analysis have enabled researchers to integrate information across biological levels, spatial scales, and disciplines. Consequently, a key difference between the earliest and most recent NoWPaS conferences is not only the increasing emphasis on adaptation, restoration, and long-term conservation, but also the growing capacity of researchers to investigate these challenges through holistic and multidisciplinary frameworks. The expansion of methodological and technological possibilities has therefore contributed to a more comprehensive understanding of Salmonid systems, allowing the scientific community to move from localized and discipline-specific questions toward increasingly integrated perspectives on ecological and conservation issues.

### 4.3 Future directions of salmonid research

#### 4.3.1 Salmonid network in the coming years

Over the past two decades, major scientific shifts were identified within the NoWPaS community, but it is not just the research presented at NoWPaS that has changed over time, so does the community. There have been marked improvements in gender parity due to the network’s proactive stance on promoting diversity and accessibility. However, there are other important metrics on community diversity that were not covered in this study, such as deprivation, disability status, and ethnic or cultural background. A comprehensive assessment of inclusion would require further investigation and data collection but should be considered in the organization of future meetings. Our results also demonstrate that around 50% of the NoWPaS community leaves academia for other roles within 10 years of presenting at the conference. This demonstrates that ideas, research and experiences shared at NoWPaS not only facilitate academic pursuits, but they likely go onto inform management, governance, policy and industry in the long-term. The network could capitalize on this by including more of these themes in the conference proceedings as well as by creating a more cohesive community with previous NoWPaS alumni.

In the coming years, the network is likely to continue expanding as awareness of NoWPaS spreads across the Northern Hemisphere. This expansion is already reflected in the growth of the community itself, but also in the increasing diversity of scientific knowledge exchanged within it. For instance, we identified a marked increase in the prevalence of studies on Arctic char in recent years, which may be partly associated with the growing representation of *Genetics and genomics*, given their good representation in this category, with Arctic char being a valuable model species for investigating eco-evolutionary processes. In this regard, diversification within the NoWPaS community does not merely bring more researchers to the table; it also broadens discussions and exchanges to encompass a wider range of species, ecosystems, and scientific questions.

In this context, we propose two complementary avenues for the future development of the network. First, given that NoWPaS is a relatively small conference designed to foster scientific exchange and networking, a similar event could be established for the Pacific community. A Pacific-focused ECR meeting would provide an opportunity to showcase salmonid research conducted in western Canada and the United States, South America, and eastern Asia, which has been largely absent from previous NoWPaS events. Second, a larger-scale salmonid conference could be considered in the coming years (as a complement to *Advances in the Population Ecology of Stream Salmonids*, which takes place every four years but does not cover coastal and marine ecosystems, a gap we identify as critical in this review). Such an event could occupy an intermediate position between large scientific societies and their associated conferences (*e.g*., FSBI, AFS, and DiadES) and smaller, more specialized meetings such as NoWPaS. A global salmonid conference could promote international research collaboration and interdisciplinary projects between Atlantic and Pacific research communities, while facilitating coordinated efforts to address the two major drivers identified in this study: *Aquaculture* and *Climate change*. Both pressures are likely to intensify in the coming years, increasing risks to wild populations and potentially influencing research priorities.

Recent studies highlight the need not only to improve our understanding of salmonid responses to rapidly changing environmental conditions, but also to develop and implement evidence-based management interventions capable of sustaining wild populations (Palm et al., 2007; Nyqvist et al., 2017, 2018; Larranaga et al., 2019). Achieving this objective requires balancing long-term knowledge acquisition with immediate conservation needs, thereby fostering integrated research frameworks that advance both ecological understanding and concrete conservation outcomes. In this light, advances in salmonid science will likely continue to unite fundamental and applied research, with researchers and managers needing to combine both strands to better grasp salmonids’ adaptive potential in the face of rapid environmental change. While some fields have achieved this, it appears that others, such as *Evolutionary Biology* or *Physiology*, are not being as frequently applied to real world problems, demonstrating a limiting theoretical gap and area for future innovation. Alternatively, it is possible that research from some countries, like the UK, could continue to become more applied, to the detriment of fundamental bluesky research. The current limits and possible perspectives to enhance the transfer from the theory to management are discussed in the next section.

#### 4.3.2 A prospect for salmonid science

Far more studies were conducted in freshwater when compared to estuarine and marine ecosystems. We expect more work in the future to be conducted in marine environments as tracking technologies and methodologies advance, as is already happening (Willmes et al., 2025; Elliott et al., 2026; Lennox et al., 2026). Satellite and acoustic telemetry, for example, are increasingly making it feasible to follow anadromous salmonids though their marine journey (Kristensen et al., 2019; Matley et al., 2022). We see this as an important frontier: future research would ideally achieve more balanced coverage across the full life-history continuum of each species. A more specific illustration of this gap is the limited research on Atlantic salmon kelts and carcasses (but see Perry et al., 2026); recent interest in kelt reconditioning as a population recovery strategy and in the ecological connectivity provided by carcasses-derived nutrients transfer (McLennan et al., 2019; Bernthal et al., 2022) may become an increasingly active area of research. This example further highlights the value of salmonids as model organisms for investigating complex ecological processes, including functional connectivity (Sturrock et al., 2026b).

The methodological trends identified in this review also highlight several promising developments. The continued diversification of molecular tools suggests that future research is well positioned to move beyond characterizing genetic diversity and toward understanding its functional expression and adaptive significance, particularly as sequencing costs continue to fall and multi-omics approaches become more accessible. This trajectory echoes a recent assessment of early predictions made during the genomics revolution, which highlighted salmonids as frontrunners in translating genomic tools into applied outcomes, while also identifying which of these predictions have yet to be fully realized (May et al., 2025). However, applying these methods more broadly, beyond Arctic charr and Atlantic salmon, is another area requiring progress. The application of high-throughput sequencing to eDNA workflows is also opening another frontier to genomic applications, being used for species monitoring in freshwater environments, that would have previously been performed using techniques such as electrofishing, now allowing catchment-scale monitoring of salmonids.

Advances in the use of archival tags similarly hold promise for improving our understanding of salmonid life history, especially in environments that have historically been difficult to study, such as the marine phase of migratory forms. For instance, biochronological archives (*e.g.*, scales, otolith, eye lenses) are increasingly used as valuable natural proxies for improving age estimation (Limburg et al., 2018; Heimbrand et al., 2020; Fontaine et al., 2024), reconstructing individual life-history trajectories (Chat et al., 2022; Reis-Santos et al., 2022; Källo et al., 2023), or investigating ecophysiological responses to environmental variability (Bareille et al., 2024; Hüssy et al., 2024). The complementary information preserved in different archival structures has also stimulated the integration of multiple chemical tracers within the same study. For example, combining otolith and eye lens chemistry has recently provided new insights into the relative contributions and ecological responses of distinct phenotypes under ongoing climate change (Sturrock et al., 2026a).

Acoustic telemetry has also shown increasing popularity in recent years due to technological advancements and associated novel insights, and this is likely to continue. However, these technologies do rely on acoustic arrays, limiting their application. Satellite tags do not have this limitation, but their high cost and large size appear to have suppressed their application among NoWPaS researchers, with no delegate talks having used this technology. Satellite tags do produce lower spatiotemporal resolution data when compared to acoustic telemetry, but they can also collect environmental data (Ismail et al., 2026) and have been used to high effect by the broader salmonid community, especially work on the marine migration of Atlantic salmon (Lacroix, 2013; Rikardsen et al., 2021). Improvements in the miniaturisation and reducing costs could therefore make satellite tagging a crucial tool for plugging the marine research gap.

Together, these methodological improvements offer the chance of more nuanced understanding that goes beyond dogmas based on traditional traits like growth, survival and movement. Expanding these tools to a broader range of species beyond Atlantic salmon and brown trout would strengthen our understanding of species resilience to environmental changes across the salmonid family. More broadly, as genetic and archival methods continue to grow, we anticipate growing opportunities to link molecular mechanisms to life history traits. Realizing this potential, however, will also require a stronger applied focus: translating these advances into management-relevant outputs, such as informing breeding and conservation programmes, remains an important direction for future NoWPaS research (Betts et al., 2026a). The continuing understanding of how habitats develop, assemble, diversify, and connect to each other across spatial and temporal scales must remain a priority in the salmonid research agenda to maintain, restore and conserve wild populations. In this regard, research has long shown the importance of habitat heterogeneity and lateral/longitudinal connectivity for salmonid populations, and when these dimensions are taken into account in practice they lead to successful small- to large-scale restoration projects (*e.g*., Munsch et al., 2023; Cordoleani et al., 2024; Roni et al., 2025; Goodman et al., 2026). While restoration of habitat diversity or connectivity focuses on degraded ecosystems, research is also bringing to the table the importance of looking at regions where new environments are becoming suitable to salmonids in response to global change (*e.g*., Pitman et al., 2021; Poulin et al., 2025; Roncoroni, 2024; Saboret et al., 2025). This should not be seen as a positive outcome of global change, but a result that merits further investigation as these environments might provide suitable conditions to salmonids in the coming decades and thus require prompt protection from harmful human activities.

Finally, we ask whether current research effort, concentrated heavily on Atlantic salmon and brown trout, may be inhibiting the ecological and management understanding of other salmonids, if research effort tracks economic and cultural value rather than ecological importance or vulnerability. A promising counterpoint is the consistent appearance of Arctic char in more recent years. Identifying the levers enabling such shifts, whether increased conservation concern, climate-driven change, or emerging funding opportunities, could help inform strategies to broaden research effort more equitably across the salmonid family in the future.

#### 4.3.3 Future roadmap

We highlight in section 4.3.2 the overarching prospects for future salmonids research, with a few choice examples. However, the data extracted in this study allows broader analyses to be conducted which give a more granular view of understudied and well-studied research areas in the NoWPaS network. Specifically, the heatmaps in Figure S3 show combinations of species with habitats, life stages, disciplines, methods, traits and drivers that are understudied or well-studied. However, in addition to this, we have produced an interactive co-occurrence plot, which shows under-represented and over-represented links between different study characteristics (Figure 9, and see Figure S10 for all links). This is a valuable roadmap for prospective future research, identifying potential research and evidence gaps. Some of the under-represented links are to be expected, for example, river-aquaculture (given that most aquaculture occurs in coastal environments or on land-based facilities), behaviour-eggs (given that eggs themselves are not able to show behaviours) or molecular approaches-movement ecology (given that there are more natural methodologies for understanding movement than molecular approaches). Some of the well-studied links are also to be expected, for example, smolt-timing, aquaculture-disease and parasitology, and fecundity-adult. While these may seem obvious, the blatancy provides a level of validation of this approach for looking at understudied and well-studied research areas. There are then the no so obvious under-represented links which could inform novel and impactful research questions, such as climate change-movement ecology, experimental approach-adult diet and management restoration-physiology, to name just a few. While the figure 9 shows under-represented links, it should be highlighted that it does not visualise links that have never been seen before at NoWPaS, and therefore truly novel combinations are absent, but this property of the network can also help filter out nonsense or infeasible combinations.

**Figure 9:**
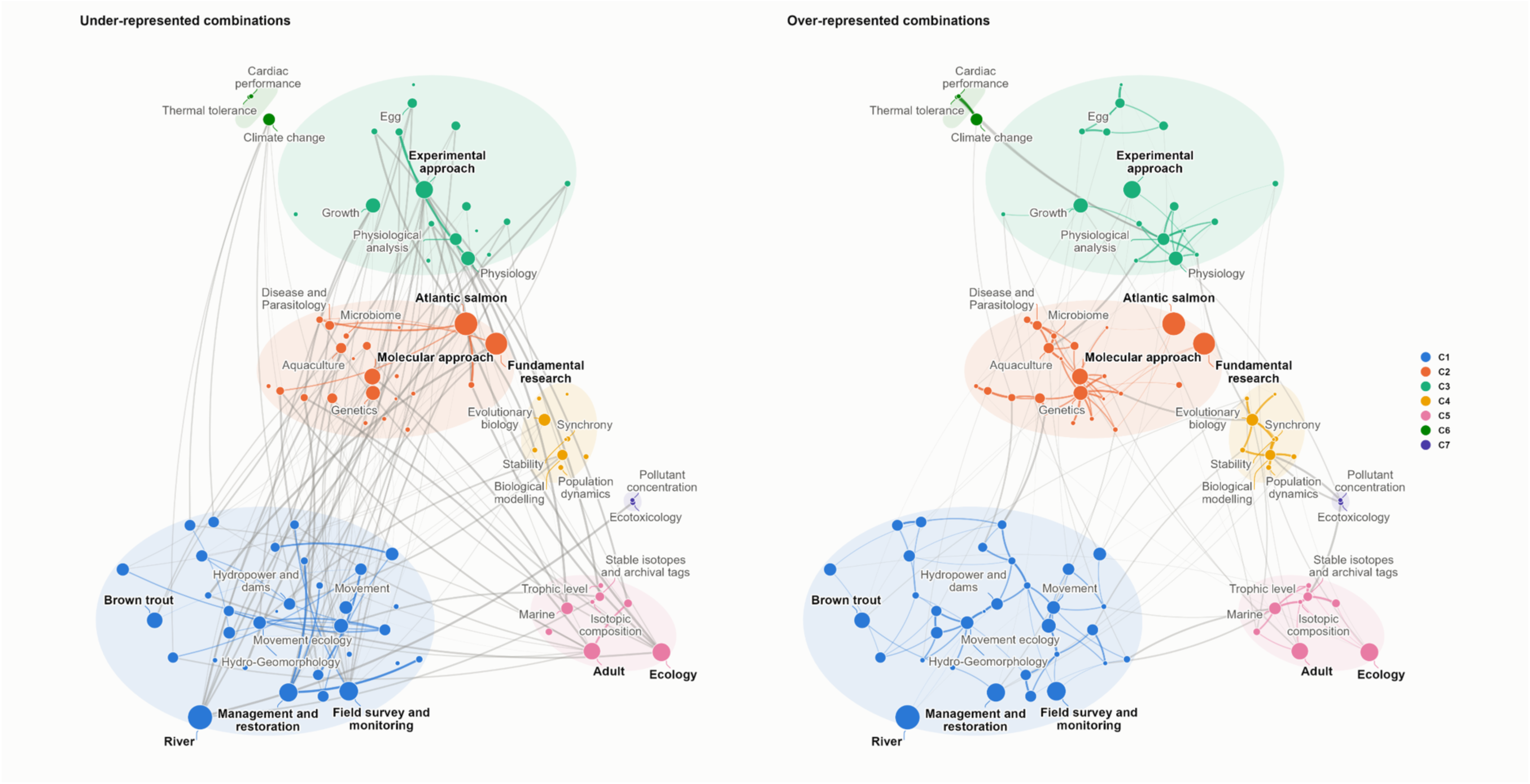
Co-occurrence network resulting from the 555 talks presented at the NoWPaS meetings between 2005 and 2025. Nodes are the categories of the seven coded variables (discipline, method, species, life stage, habitat, trait and driver) and node area is proportional to the number of talks featuring the category. Links connect pairs of categories co-occurring in at least three talks and are weighted by their association strength, *i.e*., the ratio of observed to expected co-occurrences given the frequencies of the two categories (Van Eck and Waltman, 2010). Left panel: Under-represented combinations; all links with an observed/expected ratio < 1. Right panel: Over-represented combinations; all links with an observed/expected ratio > 1. Ellipses identify the seven clusters (C1–C7) obtained with the Louvain algorithm on the entire network (200 runs, Q = 0.42). Labels show the five most central categories of each cluster and, in bold, the ten most frequent categories overall. An interactive version of this figure with the list of under- and over-represented combinations is available at https://alamarins.github.io/NoWPaS_review/NoWPaS_network_interactive_light.html.

The clustering of study characteristics also provides a useful visualisation of thematic silos which naturally fall together. Mixing study characteristics between these thematic silos is an approach for generating potentially innovative and impactful research designs and questions. For example, mixing the more molecular-disease (orange) cluster with the experimental approach-physiology (light green) cluster. Indeed, smaller clusters may also provide fruitful ground for future development and interdisciplinarity, such as the climate change (dark green) cluster or ecotoxicology (purple) cluster.

## 5 ACKNOWLEDGMENTS

We would like to thank all of the attendees, committee members and keynote speakers who have contributed to NoWPaS, as well as John Conallin, Amy Ellison and Edel Lheureux for providing abstract books. We would also like to thank the following funders who have supported NoWPaS through the years: Atlantic Salmon Trust, The FSBI, The Company of Biologists, Statkraft, Ocean Tracking Network, Research Council of Norway, INRAE, Innovasea, Norwegian University of Science and Technology, Karlstad University, Université de Pau et des Pays de l’Adour, Marine Alliance for Science and Technology for Scotland, Canadian Journal of Fisheries and Aquatic Science, Biomark, Wild Trout Trust, Thelma Biotel, PyroScience, SINTEF Energy Research, Roskilde University, Villaksprogrammet, Statkraft Energy AS, Statkraft Development AS, The Norwegian Directorate of Nature, Sintef Energy Research AS, The University of Jyväskylä, Elforsk, The Nordic Marine Academy, NNCOLD, France Hydro Électricité, Inland Fisheries Ireland, Vemco, Saumon Québec, Big Fin Scientific, Marine Scotland, The Fishmongers’ Company, Office français de la biodiversité, Fédération Nationale de la Pêche en France, Wessex Salmon and Rivers Trust, Ecole doctorale sciences exactes et leurs applications, Ministère de l’Enseignement supérieur et de la Recherche, Aquitaine, Agence de l’eau Adour-Garonne, Ecole doctorale SIBAGHE, CEDREN, Havforskningsinstituttet, International Commission on Large Dams, Dŵr Cymru Welsh Water, Environment Agency Wales, Llandysul Angling Association, Formas, Sveriges lantbruksuniversitet (SLU), Oregon RFID, Fortum, Interreg Sverige-Norge, Kungliga Skogs-och Lantbruksakademien, Vattenfall, CIRSA, Institut national de la recherche scientifique, Hydro-Québec, Fédération québécoise pour le saumon atlantique, Smith-Root, Canadian Rivers, Institute, American Fisheries Society, Groupe de recherche interuniversitaire en limnologie, Québec ville et région, Uni Research, Springhead Trust, Naturpolis, University of Eastern Finland, Niemi-säätiö, The Genetics Society, Lotek Wireless, Loligo Systems, VAKI, Landsvirkjun, Government of Iceland, Ministry for the Environment and Natural Resources (Iceland), Star-Oddi, Landssamband veiðifélaga, E2S UPPA, SSE renewables, SULSA, Atlantic Salmon Federation, Nordic Society Oikos and Nova Scotia Salmon Association.

M.R. was supported by the Swiss National Science Foundation grants 217903 and 235364. W.B.P was supported by NERC, grant NE/X015610/1. A.F. was supported by a NSERC Alliance Funds (ALLRP 560330 – 20). L.E.H was supported by the UKRI Biotechnology and Biological Sciences Research Council (BBSRC) grant numbers BB/T00875X/1 and BB/Z515267/1. A.L. was supported by the European Union (ERC, FishLEGs, 101054307). Views and opinions expressed are however those of the author(s) only and do not necessarily reflect those of the European Union or the European Research Council Executive Agency. Neither the European Union nor the granting authority can be held responsible for them.

## 6 DATA AND CODE AVAILABILITY

The database and code used for this review are available at https://github.com/alamarins/NoWPaS_review.

