## Supplemental File for "SALMONID SCIENCE ON THE MOVE: TWENTY YEARS OF RESEARCH THROUGH THE LENS OF EARLY CAREER RESEARCHERS"

### SUPPLEMENTARY TABLES

Table S1: Research disciplines investigated in this study along with the corresponding sub-disciplines or types of studies.

| Research discipline | Sub-discipline / Studies type |
|---|---|
| Hydro-geomorphology | Eco-hydraulics, hydrology, fluvial geomorphology, hyporheic science, hydropower fishways, habitat suitability criteria, physical habitat modelling, remote sensing |
| Behavioural ecology | |
| Movement ecology | Hydropower fishway passage, migration studies |
| Habitat use | |
| Ecology | Population dynamics, studies of the effect of the environment on life histories, of density / competition, food webs, ecosystems |
| Evolutionary biology | Life history evolution, sexual selection, GxE interaction, plasticity |
| Physiology | Endocrinology, immunology, ecophysiology |
| Ecotoxicology | |
| Disease and Parasitology | Epidemiology, parasitology, disease ecology |
| Genetics and Genomics | Population Genetics, phylogeography/ phylogeny, gene expression, epigenetics, genomics |

Table S2: Research methods investigated in this study.

| **Research method** | **Examples** |
|---|---|
| Experimental approach | Channel, flume, enrichment, behavioural, common-garden, field, lab experiments |
| Physiological analysis | Respirometry, lipid analysis, mass spectrometry, metabolomics, histology, stress analysis, gametes analysis |
| Stable isotopes and archival tags | Stable isotopes, chemistry of natural tags (otolith, scale, eye lens) |
| Field survey and monitoring | In situ monitoring (electrofishing, snorkeling, video monitoring, mark-recapture), telemetry, parasite infection rate, morphology, scalimetry, stomach content |
| Molecular approach | Gene expression, allozyme, epigenomics, microsatellites, GWAS, eDNA, Genomic offset, Proteomics |
| Biological modelling | Bioenergetic modelling, population modelling, simulation model |
| Habitat characterization | Habitat modelling, hydraulic modelling, GIS, remote sensing |
| Literature review | - |

Table S3: Total occurrences of every genus (or group) and species (or group), both for Salmonid and non-Salmonid species.

| Genus/Group | Species/Group | Scientific name | Count |
|---|---|---|---|
| | Salmonid species | | |
| *Salmo* | Atlantic salmon | *S. salar* | 329 |
| | Atlantic salmon (landlocked) | *S. salar* | 17 |
| | Brown trout | *S. trutta* | 136 |
| | Sea trout | *S. trutta* | 57 |
| | Marble trout | *S. marmoratus* | 3 |
| | *Total* | | *542* |
| *Salvelinus* | Arctic char | *S. alpinus* | 18 |
| | Arctic char (anadromous) | *S. alpinus* | 11 |
| | Brook char | *S. fontinalis* | 13 |
| | Brook char (anadromous) | *S. fontinalis* | 2 |
| | Dolly Varden | *S. malma* | 4 |
| | Lake char | *S. umbla* | 1 |
| | Lake trout | *S. namaycush* | 1 |
| | *Total* | | *50* |
| *Oncorhynchus* | Rainbow trout | *O. mykiss* | 7 |
| | Coho salmon | *O. kisutch* | 4 |
| | Chinook salmon | *O. tshawytscha* | 1 |
| | Pink salmon | *O. gorbuscha* | 1 |
| | *Total* | | *13* |
| *Coregonus* | Atlantic whitefish | *C. huntsmani* | 1 |
| | European whitefish | *C. lavaretus* | 1 |
| | Whitefish (species not specified) | | 1 |
| | *Total* | | *3* |
| *Thymallus* | European grayling | *T. thymallus* | 3 |
| | *Total* | | *3* |
| *NA* | Salmonid (species not specified) | | 17 |
| | *Total* | | *17* |
| | Non-Salmonid species | | |
| Freshwater fish | Bullhead | | 3 |
| | Burbot | | 3 |
| | Northern pike | *Esox lucius* | 2 |
| | European eel | *Anguilla anguilla* | 1 |
| | Lamprey | | 1 |
| | Fallfish | *Semotilus corporalis* | 1 |
| | *Galaxiids* | | 1 |

| | | | |
|---|---|---|---|
| | Freshwater fish (species not specified) | | 1 |
| | *Total* | | *13* |
| Saltwater fish | European flounder | *Platichthys flesus* | 2 |
| | Atlantic cod | *Gadus morhua* | 1 |
| | Herring | | 1 |
| | Saithe | | 1 |
| | Sprat | | 1 |
| | Striped bass | *Morone saxatilis* | 1 |
| | *Total* | | *7* |
| Parasite | Pearl mussel | | 5 |
| | Tetracapsuloides bryosalmonae | | 5 |
| | *Gyrodactylus* | | 3 |
| | Sea lice | | 3 |
| | Freshwater fish lice | | 1 |
| | Parasite (not specified) | | 2 |
| | *Total* | | *19* |
| Freshwater organism | Didymo | | 7 |
| | *Daphnia magna* | | 1 |
| | Macrozoobenthos | | 1 |
| | *Total* | | *9* |
| Terrestrial vegetation | Japanese knotweed | | 1 |
| | *Total* | | *1* |
| Aquatic vegetation | Macrophyte | | 4 |
| | *Total* | | *4* |
| Mammal | Eurasian beaver | *Castor fiber* | 5 |
| | Otter | | 2 |
| | *Total* | | *7* |
| Gut microbiome | Gut microbiome | | 3 |
| | *Total* | | *3* |
| NA | NA | | 9 |
| | *Total* | | *9* |

Table S4: Temporal trend in the proportion of abstracts by discipline, estimated by a quasibinomial logistic model. Estimates (log-odds change per scaled year), standard errors, test statistics, and *p*-values are rounded to 3 digits and shown for the scaled year term, fitted separately for each discipline.

| **Discipline** | **Estimate** | **SE** | **Statistic (z)** | ***p*-val** | **$p_{adj}$ (BH)** |
|---|---|---|---|---|---|
| Behavioural ecology | -0.015 | 0.130 | -0.117 | 0.908 | 0.912 |
| Disease and parasitology | 0.315 | 0.230 | 1.368 | 0.187 | 0.337 |
| Ecology | -0.01 | 0.091 | -0.112 | 0.912 | 0.912 |
| Evolutionary biology | 0.303 | 0.187 | 1.617 | 0.122 | 0.337 |
| Genetics and genomics | 0.449 | 0.108 | 4.153 | 0.001 | 0.002 |
| Habitat use | -0.191 | 0.161 | -1.187 | 0.250 | 0.375 |
| Hydro-Geomorphology | -0.686 | 0.133 | -5.16 | 0.000 | 0.001 |
| Movement ecology | 0.072 | 0.129 | 0.556 | 0.585 | 0.752 |
| Physiology | 0.106 | 0.077 | 1.375 | 0.185 | 0.337 |

***p*-val: *p*-value; $p_{adj}$ (BH): *p*-value adjusted for Benjamini-Hochberg correction; SE: Standard Error.**

Table S5: Temporal trend in the proportion of abstracts by discipline and main species (Atlantic salmon *vs.* brown and sea trout), estimated by a quasibinomial logistic model with a year x species interaction. Estimates (log-odds change per scaled year for the reference group, Atlantic salmon, and differences for brown and sea trout), standard errors, test statistics, and *p*-values are rounded to 3 digits and shown for the scaled year, species, and interaction terms, fitted separately for each discipline.

| Discipline | Term | Estimate | SE | Statistic (z) | *p*-val | $p_{adj}$ (BH) |
|---|---|---|---|---|---|---|
| **Behavioural ecology** | Year | -0.039 | 0.153 | -0.253 | 0.802 | 0.802 |
| **Behavioural ecology** | Brown trout + Sea trout | -0.362 | 0.245 | -1.481 | 0.147 | 0.33 |
| **Behavioural ecology** | Year:Brown trout + Sea trout | -0.246 | 0.237 | -1.039 | 0.306 | 0.459 |
| **Disease and parasitology** | Year | 0.454 | 0.265 | 1.714 | 0.095 | 0.284 |
| **Disease and parasitology** | Brown trout + Sea trout | -0.734 | 0.472 | -1.554 | 0.128 | 0.315 |
| **Disease and parasitology** | Year:Brown trout + Sea trout | -0.638 | 0.455 | -1.402 | 0.169 | 0.351 |
| **Ecology** | Year | 0.111 | 0.105 | 1.056 | 0.298 | 0.459 |
| **Ecology** | Brown trout + Sea trout | -0.855 | 0.186 | -4.586 | 0 | 0.001 |
| **Ecology** | Year:Brown trout + Sea trout | -0.599 | 0.18 | -3.33 | 0.002 | 0.01 |
| **Evolutionary biology** | Year | 0.344 | 0.195 | 1.76 | 0.086 | 0.284 |
| **Evolutionary biology** | Brown trout + Sea trout | -0.32 | 0.305 | -1.049 | 0.301 | 0.459 |
| **Evolutionary biology** | Year:Brown trout + Sea trout | -0.292 | 0.296 | -0.988 | 0.33 | 0.459 |
| **Genetics and genomics** | Year | 0.395 | 0.138 | 2.873 | 0.007 | 0.03 |
| **Genetics and genomics** | Brown trout + Sea trout | -1.189 | 0.271 | -4.39 | 0 | 0.001 |
| **Genetics and genomics** | Year:Brown trout + Sea trout | -0.47 | 0.263 | -1.786 | 0.082 | 0.284 |
| **Habitat use** | Year | -0.308 | 0.19 | -1.624 | 0.113 | 0.304 |
| **Habitat use** | Brown trout + Sea trout | -0.267 | 0.293 | -0.912 | 0.368 | 0.473 |
| **Habitat use** | Year:Brown trout + Sea trout | 0.186 | 0.284 | 0.654 | 0.517 | 0.635 |

| | | | | | | |
|---|---|---|---|---|---|---|
| **Hydro-geomorphology** | Year | -0.703 | 0.188 | -3.732 | 0.001 | 0.004 |
| **Hydro-geomorphology** | Brown trout + Sea trout | -0.324 | 0.322 | -1.004 | 0.321 | 0.459 |
| **Hydro-geomorphology** | Year:Brown trout + Sea trout | -0.284 | 0.294 | -0.966 | 0.34 | 0.459 |
| **Movement ecology** | Year | 0.041 | 0.142 | 0.29 | 0.773 | 0.802 |
| **Movement ecology** | Brown trout + Sea trout | -0.293 | 0.219 | -1.336 | 0.19 | 0.366 |
| **Movement ecology** | Year:Brown trout + Sea trout | -0.069 | 0.214 | -0.323 | 0.748 | 0.802 |
| **Physiology** | Year | 0.04 | 0.083 | 0.489 | 0.628 | 0.737 |
| **Physiology** | Brown trout + Sea trout | -1.002 | 0.154 | -6.499 | 0 | 0 |
| **Physiology** | Year:Brown trout + Sea trout | 0.043 | 0.15 | 0.286 | 0.777 | 0.802 |

***p*-val: *p*-value; $p_{adj}$ (BH): *p*-value adjusted for Benjamini-Hochberg correction; SE: Standard Error.**

Table S6: Temporal trend in the proportion of abstracts by method, estimated by a quasibinomial logistic model. Estimates (log-odds change per scaled year), standard errors, test statistics, and *p*-values are rounded to 3 digits and shown for the scaled year term, fitted separately for each method.

| **Method** | **Estimate** | **SE** | **Statistic (z)** | ***p*-val** | **$p_{adj}$ (BH)** |
|---|---|---|---|---|---|
| **Biological modelling** | 0.071 | 0.177 | 0.403 | 0.692 | 0.834 |
| **Experimental approach** | -0.157 | 0.103 | -1.525 | 0.144 | 0.335 |
| **Field survey and monitoring** | 0.026 | 0.121 | 0.213 | 0.834 | 0.834 |
| **Habitat characterization** | -0.37 | 0.138 | -2.685 | 0.015 | 0.051 |
| **Molecular approach** | 0.508 | 0.103 | 4.914 | 0 | 0.001 |
| **Physiological analysis** | 0.028 | 0.119 | 0.233 | 0.819 | 0.834 |
| **Stable isotopes and archival tags** | 0.167 | 0.174 | 0.965 | 0.347 | 0.607 |

***p*-val: *p*-value; $p_{adj}$ (BH): *p*-value adjusted for Benjamini-Hochberg correction; SE: Standard Error.**

Table S7: Temporal trend in the proportion of abstracts by method and main species (Atlantic salmon *vs.* brown and sea trout), estimated by a quasibinomial logistic model with a year x species interaction. Estimates (log-odds change per scaled year for the reference group, Atlantic salmon, and differences for brown and sea trout), standard errors, test statistics, and *p*-values are rounded to 3 digits and shown for the scaled year, species, and interaction terms, fitted separately for each method.

| **Method** | **Term** | **Estimate** | **SE** | **Statistic (z)** | ***p*-val** | **$p_{adj}$ (BH)** |
|---|---|---|---|---|---|---|
| **Biological modelling** | Year | -0.05 | 0.177 | -0.28 | 0.781 | 0.82 |
| **Biological modelling** | Brown trout + Sea trout | -1.005 | 0.351 | -2.865 | 0.007 | 0.024 |
| **Biological modelling** | Year:Brown trout + Sea trout | 0.45 | 0.336 | 1.337 | 0.189 | 0.284 |
| **Experimental approach** | Year | -0.106 | 0.101 | -1.049 | 0.301 | 0.421 |
| **Experimental approach** | Brown trout + Sea trout | -0.803 | 0.179 | -4.482 | 0 | 0.001 |
| **Experimental approach** | Year:Brown trout + Sea trout | -0.429 | 0.173 | -2.486 | 0.017 | 0.045 |
| **Field survey and monitoring** | Year | 0.101 | 0.127 | 0.798 | 0.43 | 0.564 |
| **Field survey and monitoring** | Brown trout + Sea trout | -0.409 | 0.199 | -2.055 | 0.047 | 0.089 |
| **Field survey and monitoring** | Year:Brown trout + Sea trout | -0.418 | 0.194 | -2.158 | 0.037 | 0.078 |
| **Habitat characterization** | Year | -0.428 | 0.175 | -2.446 | 0.019 | 0.045 |
| **Habitat characterization** | Brown trout + Sea trout | -0.438 | 0.294 | -1.489 | 0.145 | 0.253 |
| **Habitat characterization** | Year:Brown trout + Sea trout | -0.171 | 0.278 | -0.617 | 0.541 | 0.668 |
| **Molecular approach** | Year | 0.457 | 0.134 | 3.409 | 0.002 | 0.007 |
| **Molecular approach** | Brown trout + Sea trout | -1.137 | 0.253 | -4.497 | 0 | 0.001 |
| **Molecular approach** | Year:Brown trout + Sea trout | -0.635 | 0.246 | -2.585 | 0.014 | 0.041 |
| **Physiological analysis** | Year | 0.009 | 0.115 | 0.076 | 0.94 | 0.94 |
| **Physiological analysis** | Brown trout + Sea trout | -0.97 | 0.219 | -4.422 | 0 | 0.001 |
| **Physiological analysis** | Year:Brown trout + Sea trout | -0.304 | 0.212 | -1.433 | 0.16 | 0.258 |

| | | | | | | |
|---|---|---|---|---|---|---|
| **Stable isotopes and archival tags** | Year | 0.056 | 0.176 | 0.319 | 0.751 | 0.82 |
| **Stable isotopes and archival tags** | Brown trout + Sea trout | -1.61 | 0.431 | -3.737 | 0.001 | 0.003 |
| **Stable isotopes and archival tags** | Year:Brown trout + Sea trout | -0.189 | 0.42 | -0.449 | 0.656 | 0.765 |

**_p_-val: _p_-value; $p_{adj}$ (BH): _p_-value adjusted for Benjamini-Hochberg correction; SE: Standard Error.**

Table S8: Temporal trend in the proportion of abstracts by research driver, estimated by a quasibinomial logistic model. Estimates (log-odds change per scaled year), standard errors, test statistics, and *p*-values are rounded to 3 digits and shown for the scaled year term, fitted separately for each driver.

| **Driver** | **Estimate** | **SE** | **Statistic (z)** | ***p*-val** | **$p_{adj}$ (BH)** |
|---|---|---|---|---|---|
| **Aquaculture** | 0.394 | 0.128 | 3.086 | 0.006 | 0.018 |
| **Climate change** | 0.779 | 0.176 | 4.436 | 0 | 0.002 |
| **Fundamental research** | -0.026 | 0.087 | -0.3 | 0.768 | 0.805 |
| **Hydropower and dams** | 0.049 | 0.115 | 0.428 | 0.673 | 0.805 |
| **Management and restoration** | 0.227 | 0.11 | 2.059 | 0.054 | 0.107 |
| **Stocking** | -0.054 | 0.215 | -0.25 | 0.805 | 0.805 |

**_p_-val: _p_-value; $p_{adj}$ (BH): _p_-value adjusted for Benjamini-Hochberg correction; SE: Standard Error.**

Table S9: Temporal trend in the proportion of abstracts by research driver and main species (Atlantic salmon *vs.* brown and sea trout), estimated by a quasibinomial logistic model with a year x species interaction. Estimates (log-odds change per scaled year for the reference group, Atlantic salmon, and differences for brown and sea trout), standard errors, test statistics, and *p*-values are rounded to 3 digits and shown for the scaled year, species, and interaction terms, fitted separately for each driver.

| **Driver** | **Term** | **Estimate** | **SE** | **Statistic (z)** | ***p*-val** | **$p_{adj}$ (BH)** |
|---|---|---|---|---|---|---|
| **Aquaculture** | Year | 0.436 | 0.158 | 2.761 | 0.009 | 0.026 |
| **Aquaculture** | Brown trout + Sea trout | -1.474 | 0.357 | -4.127 | 0 | 0.002 |
| **Aquaculture** | Year:Brown trout + Sea trout | -0.541 | 0.347 | -1.561 | 0.127 | 0.208 |
| **Climate change** | Year | 0.468 | 0.179 | 2.614 | 0.013 | 0.033 |
| **Climate change** | Brown trout + Sea trout | -1.882 | 0.561 | -3.358 | 0.002 | 0.011 |
| **Climate change** | Year:Brown trout + Sea trout | 0.668 | 0.486 | 1.373 | 0.178 | 0.267 |
| **Fundamental research** | Year | -0.031 | 0.083 | -0.37 | 0.714 | 0.856 |
| **Fundamental research** | Brown trout + Sea trout | -0.839 | 0.139 | -6.03 | 0 | 0 |
| **Fundamental research** | Year:Brown trout + Sea trout | -0.437 | 0.135 | -3.229 | 0.003 | 0.012 |
| **Hydropower and dams** | Year | 0.069 | 0.169 | 0.409 | 0.685 | 0.856 |
| **Hydropower and dams** | Brown trout + Sea trout | -0.074 | 0.249 | -0.298 | 0.767 | 0.863 |
| **Hydropower and dams** | Year:Brown trout + Sea trout | -0.019 | 0.242 | -0.08 | 0.937 | 0.937 |
| **Management and restoration** | Year | 0.242 | 0.114 | 2.125 | 0.04 | 0.09 |
| **Management and restoration** | Brown trout + Sea trout | -0.542 | 0.182 | -2.987 | 0.005 | 0.018 |
| **Management and restoration** | Year:Brown trout + Sea trout | -0.31 | 0.177 | -1.748 | 0.089 | 0.177 |
| **Stocking** | Year | 0.055 | 0.273 | 0.2 | 0.843 | 0.892 |
| **Stocking** | Brown trout + Sea trout | -0.824 | 0.528 | -1.561 | 0.127 | 0.208 |

| | | | | | | |
|---|---|---|---|---|---|---|
| **Stocking** | Year:Brown trout + Sea trout | -0.645 | 0.498 | -1.293 | 0.204 | 0.282 |

***p*-val: *p*-value; $p_{adj}$ (BH): *p*-value adjusted for Benjamini-Hochberg correction; SE: Standard Error.**

Table S10: List of traits with occurrences in assignments in parentheses.

Growth (111), Movement (84), Survival (83), Behaviour (53), Genetic diversity (52), Density (49), Habitat use (49), Habitat (48), Metabolism (30), Gene expression (24), Diet (23), Morphology (22), Timing (18), Age at maturity (14), Infection rate (14), Stress (14), Introgression (12), Straying (10), Body composition (9), Reproductive success (9), Microbiome (8), Life-history trait (6), Population dynamics (6), Population size (6), Synchrony (6), Trophic level (6), Stock assignment (5), Cardiac performance (4), DNA methylation (4), Effective population size (4), Fecundity (4), Genetic architecture (4), Immune response (4), Isotopic composition (4), Mate choice (4), Pollutant concentration (4), Sperm trait (4), Swimming capacity (4), Telomere length (4), Thermal tolerance (4), Egg trait (3), Flow velocity (3), Genotype-phenotype association (3), Heritability (3), Lipid composition (3), Resistance (3), Stability (3), Adaptation (2), Age at migration (2), Allele frequency (2), Body condition (2), Carrying capacity (2), Color (2), Disease transmission (2), Fitness (2), Inbreeding (2), Osmoregulation capacity (2), Phylogeny (2), Portfolio effect (2), Swimming behaviour (2), Thermal refugia (2), Activity (1), Age at reproduction (1), Age at smoltification (1), Aggregation (1), Biodiversity (1), Biofouling (1), Biomass (1), Cage deformation (1), Cell pluripotency (1), DNA damage (1), Eutrophication (1), Evolution (1), Evolutionary rescue (1), Fertilisation rate (1), Fishing effort (1), Food intake (1), Gamete quality (1), Gene-environment association (1), Genomic reaction norm (1), Hyporheic exchange (1), Imputation accuracy (1), Invasion (1), Local adaptation (1), Maturation (1), Migration ability (1), Migration rate (1), Migration speed (1), Mitochondrial efficiency (1), Ovarian fluid viscosity (1), Parasite presence (1), Parasite survival and infectivity (1), Plasticity (1), Recapture rate (1), Reproductive activity (1), Selection signature (1), Size at maturation (1), Swimming activity (1), Vaccine efficiency (1)

Table S11: Temporal variation in Shannon diversity (*H*), lexical richness (S) and evenness (J) calculated from tokenized abstract research questions. Statistical results are presented for all disciplines combined and by research discipline.

| **All disciplines** | | | | | | |
|---|---|---|---|---|---|---|
| | **Metric** | **Estimate** | **SE** | **Statistic** | ***p*-val** | **$p_{adj}$ (BH)** |
| | H | 0.028 | 0.013 | 2.188 | 0.041 | 0.068 |
| | S | 9.445 | 4.407 | 2.143 | 0.045 | 0.068 |
| | J | 0 | 0 | -0.524 | 0.607 | 0.608 |
| **By discipline** | | | | | | |
| | **Metric** | **Slope** | **$r^2$** | **AIC** | ***p*-val** | **$p_{adj}$ (BH)** |
| Behavioural ecology | H | 0.024 | 0.048 | 44.3 | 0.351 | 0.527 |
| Disease and parasitology | H | 0.091 | 0.51 | 31.3 | 0.004 | 0.006 |
| Ecology | H | 0.033 | 0.116 | 40.2 | 0.131 | 0.196 |
| Evolutionary biology | H | 0.071 | 0.293 | 43.4 | 0.02 | 0.061 |
| Genetics and genomics | H | 0.091 | 0.538 | 34.9 | 0 | 0.001 |
| Habitat use | H | 0.017 | 0.024 | 45.6 | 0.513 | 0.513 |
| Hydro-geomorphology | H | -0.041 | 0.148 | 40.0 | 0.104 | 0.104 |
| Movement ecology | H | 0.044 | 0.192 | 40.2 | 0.047 | 0.088 |
| Physiology | H | 0.039 | 0.145 | 41.8 | 0.088 | 0.143 |
| Behavioural ecology | S | 0.961 | 0.062 | 187 | 0.288 | 0.527 |
| Disease and parasitology | S | 3.39 | 0.533 | 132 | 0.003 | 0.006 |
| Ecology | S | 3.65 | 0.119 | 238 | 0.125 | 0.196 |
| Evolutionary biology | S | 3.39 | 0.236 | 188 | 0.041 | 0.061 |
| Genetics and genomics | S | 6.08 | 0.377 | 216 | 0.004 | 0.006 |
| Habitat use | S | 1.25 | 0.074 | 195 | 0.245 | 0.513 |
| Hydro-geomorphology | S | -1.83 | 0.209 | 176 | 0.049 | 0.073 |
| Movement ecology | S | 3.23 | 0.176 | 223 | 0.059 | 0.088 |
| Physiology | S | 2.71 | 0.139 | 221 | 0.096 | 0.143 |
| Behavioural ecology | J | 0 | 0.012 | -115 | 0.649 | 0.649 |
| Disease and parasitology | J | -0.001 | 0.064 | -73.6 | 0.381 | 0.381 |
| Ecology | J | 0 | 0.001 | -124 | 0.875 | 0.875 |

| | | | | | | |
|---|---|---|---|---|---|---|
| Evolutionary biology | J | 0 | 0.008 | -109 | 0.725 | 0.725 |
| Genetics and genomics | J | 0 | 0.02 | -113 | 0.55 | 0.55 |
| Habitat use | J | 0 | 0.035 | -128 | 0.429 | 0.513 |
| Hydro-geomorphology | J | 0.001 | 0.21 | -118 | 0.048 | 0.073 |
| Movement ecology | J | 0 | 0.006 | -117 | 0.744 | 0.744 |
| Physiology | J | 0 | 0.011 | -132 | 0.646 | 0.646 |

**AIC: Akaike Information Criterion; J: Lexical evenness; H: Shannon diversity; *p*-val: *p*-value; $p_{adj}$ (BH): *p*-value adjusted for Benjamini-Hochberg correction; S: Lexical richness; SE: Standard Error.**

### SUPPLEMENTARY FIGURES

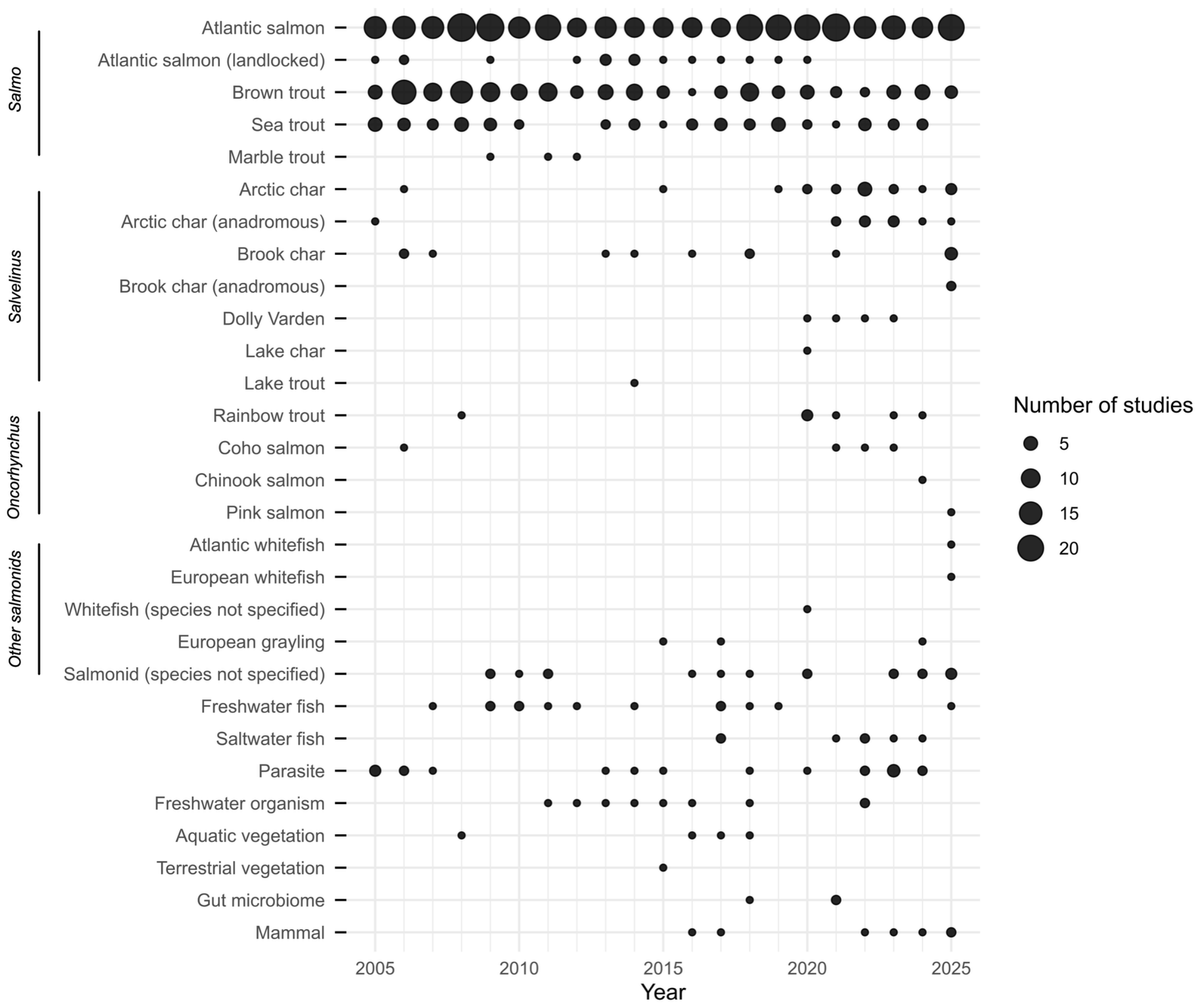


Figure S1: Annual total occurrences of every genus (or group) and species (or group), for both Salmonid and non-Salmonid species.

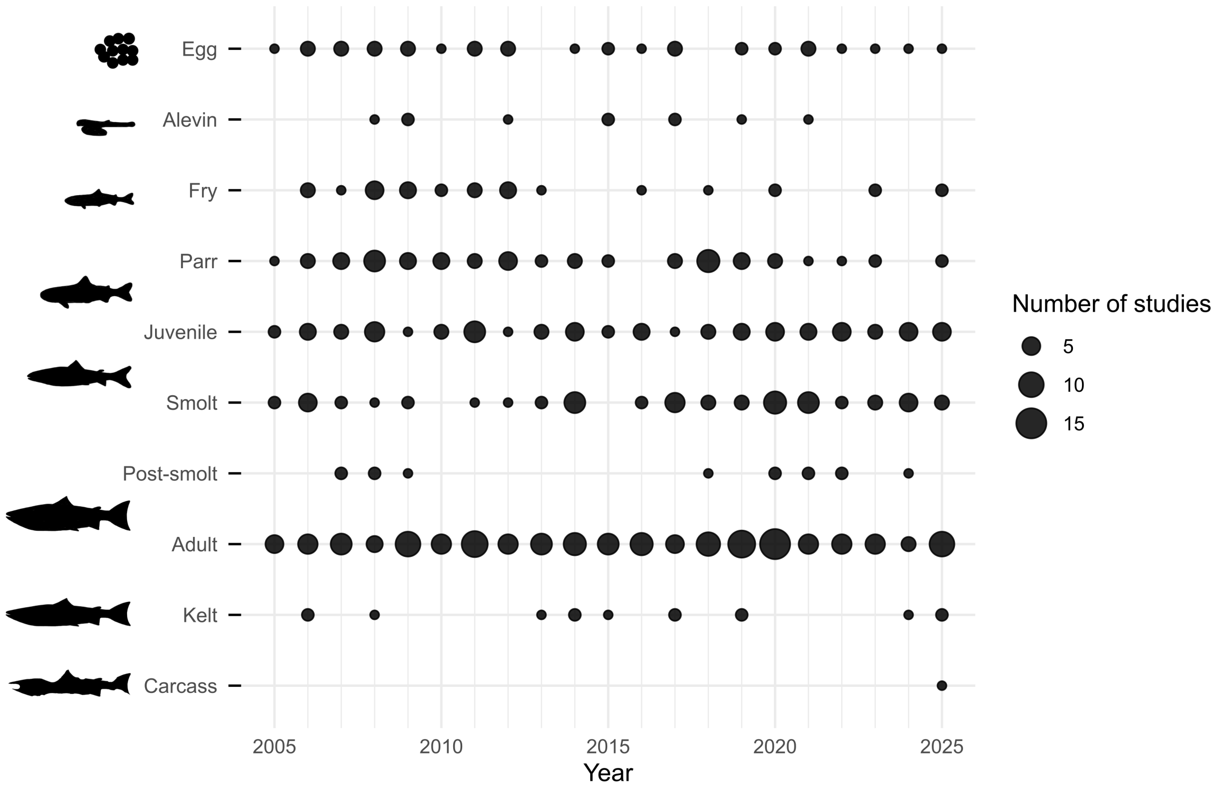


Figure S2: Lifestages studied within the NoWPaS network between 2005 and 2025. Total counts were as follows: egg, n = 38; alevin, n = 10; fry, n = 31; parr, n = 62; juvenile, n = 76; smolt, n = 65; post-smolt, n = 13; adult, n = 157; kelt, n = 14; carcass, n = 1; not specified, n = 172.

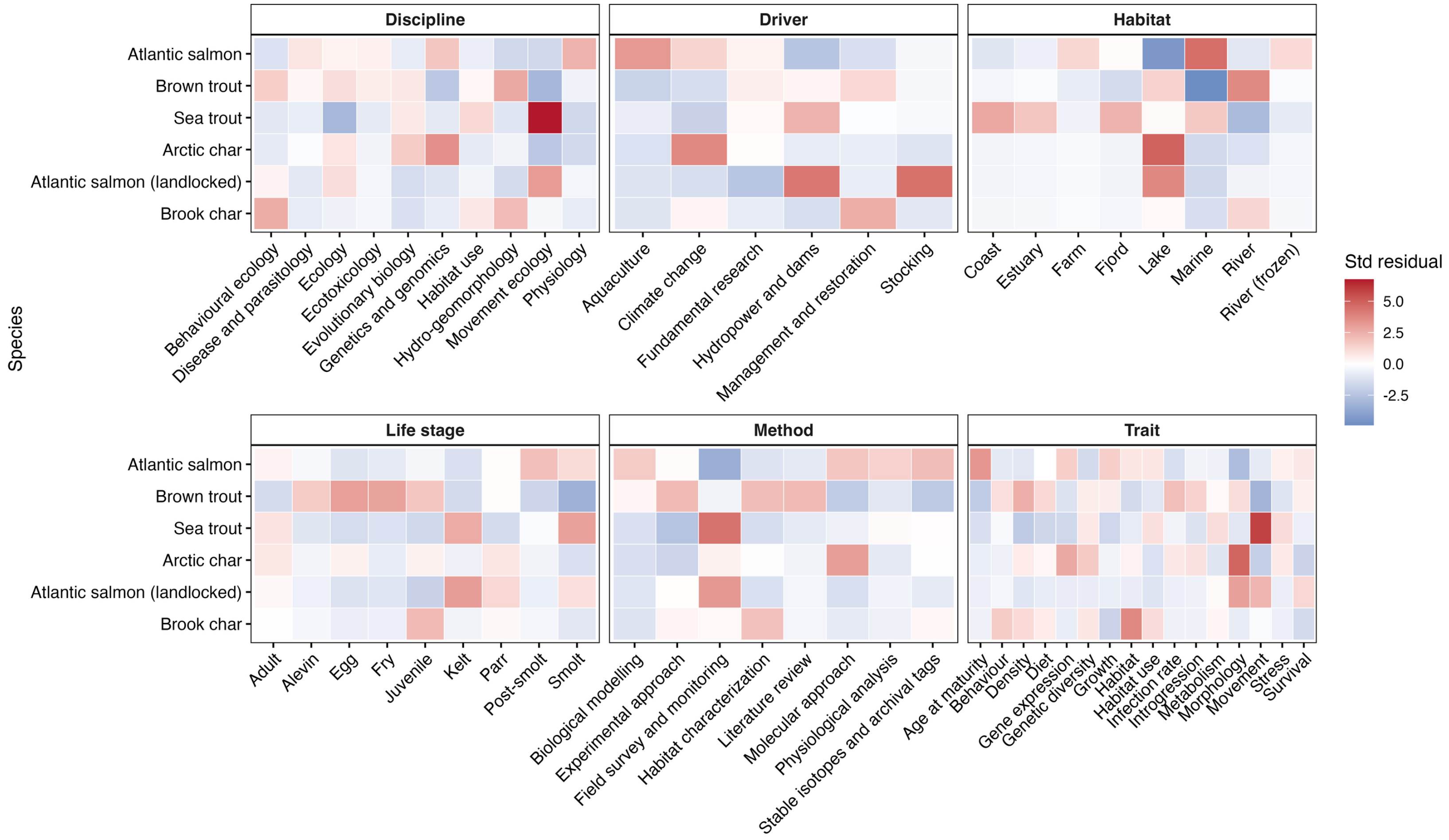


106

107 Figure S3: Co-occurrence of study species and disciplines, drivers, habitats, life stages, methods and traits measured by standardized
108 Pearson residuals from chi-squared tests of independence. Positive residuals (red) indicate more frequent combinations than expected
109 under independence, while negative residuals (blue) indicate under-represented combinations during the 2005-2025 period. Std
110 residuals: Standardized Pearson residuals.

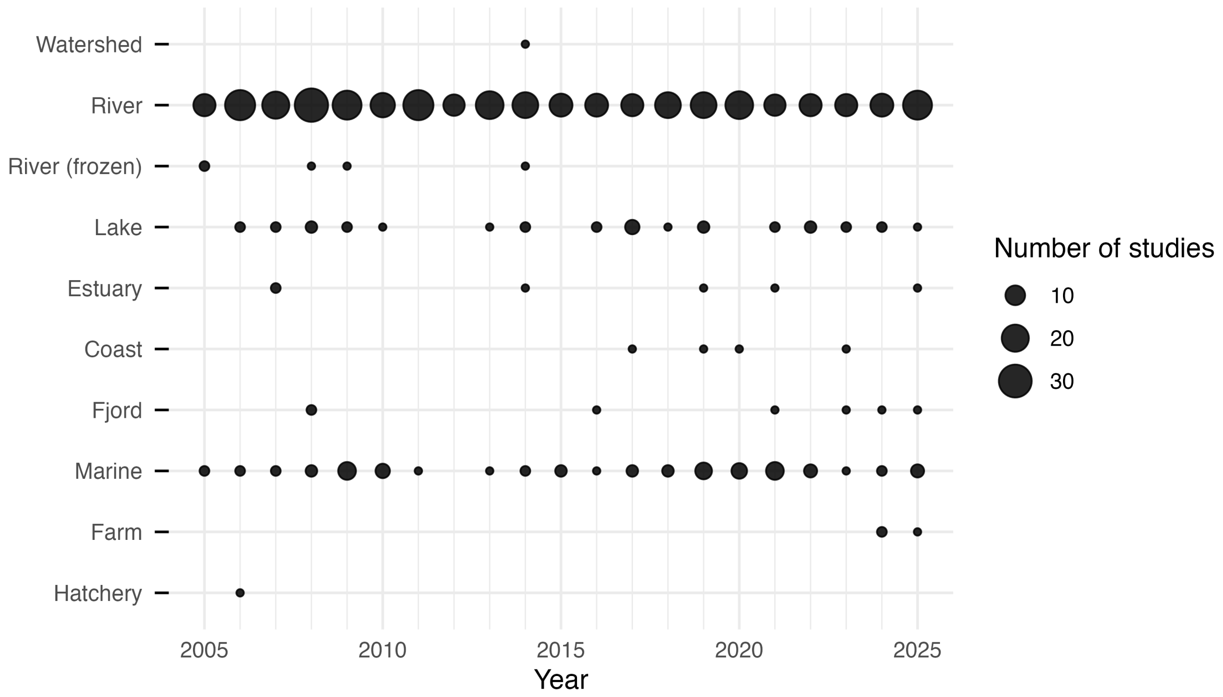


Figure S4: Environments and habitat investigated within the NoWPaS network between 2005 and 2025. Total counts were as follows: watershed, n = 1; river, n = 377; river (frozen), n = 5, lake, n = 34; estuary, n = 6; coast, n = 4; fjord, n = 7; marine, n = 68; farm, n = 3; hatchery, n = 1; not specified, n = 81.

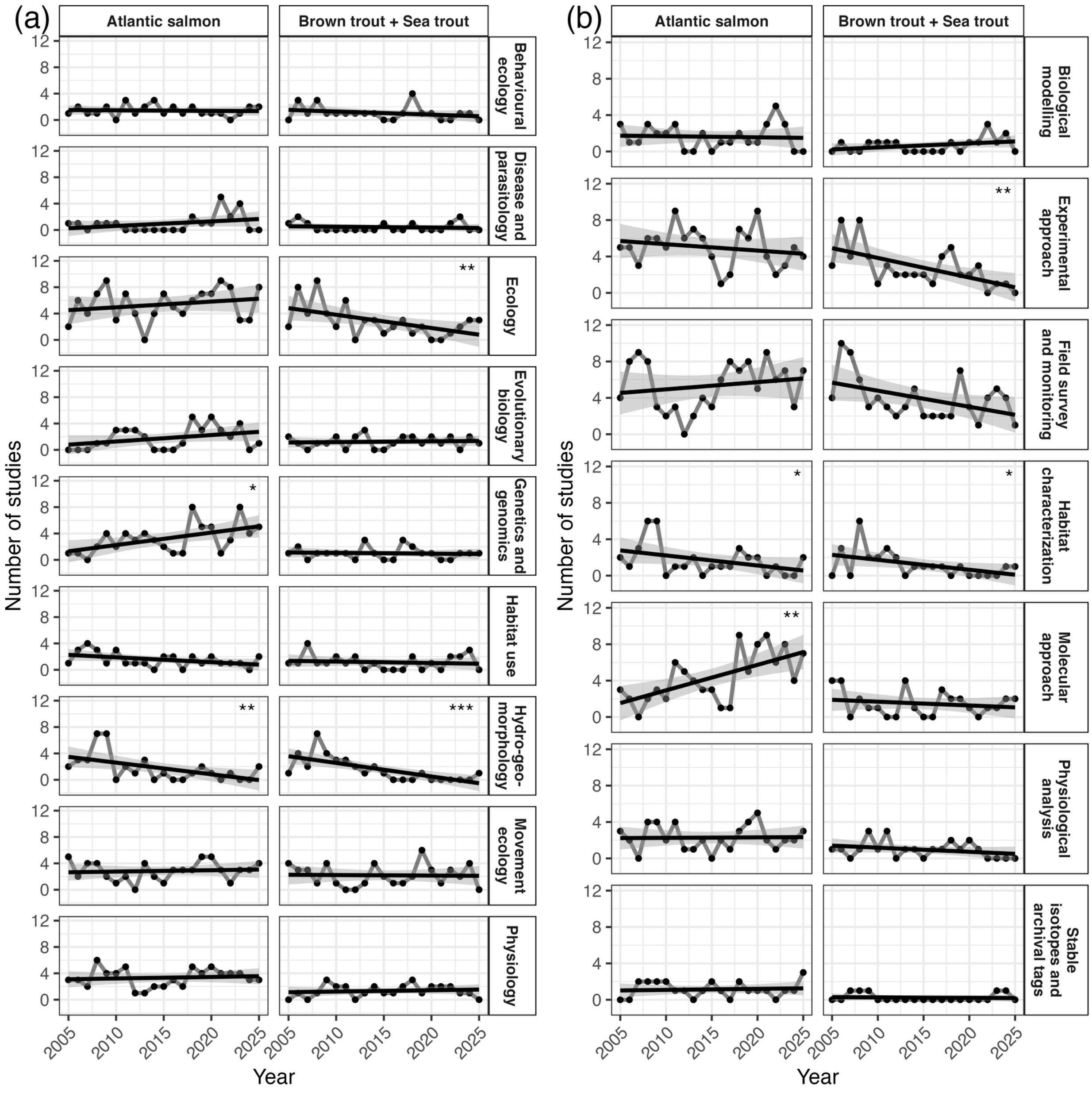


Figure S5: Temporal change in the occurrence of (a) research disciplines and (b) methods for studies involving Atlantic salmon and Brown trout and Sea trout. Points represent the annual number of studies per category. Black solid lines show linear regression fits for each category, with shaded areas indicating 95% confidence intervals around the fitted relationships. Asterisks denote statistically significant temporal trends based on the slope of the linear regression ($p <$ 0.05*, $p < 0.01$**, $p < 0.001$***).

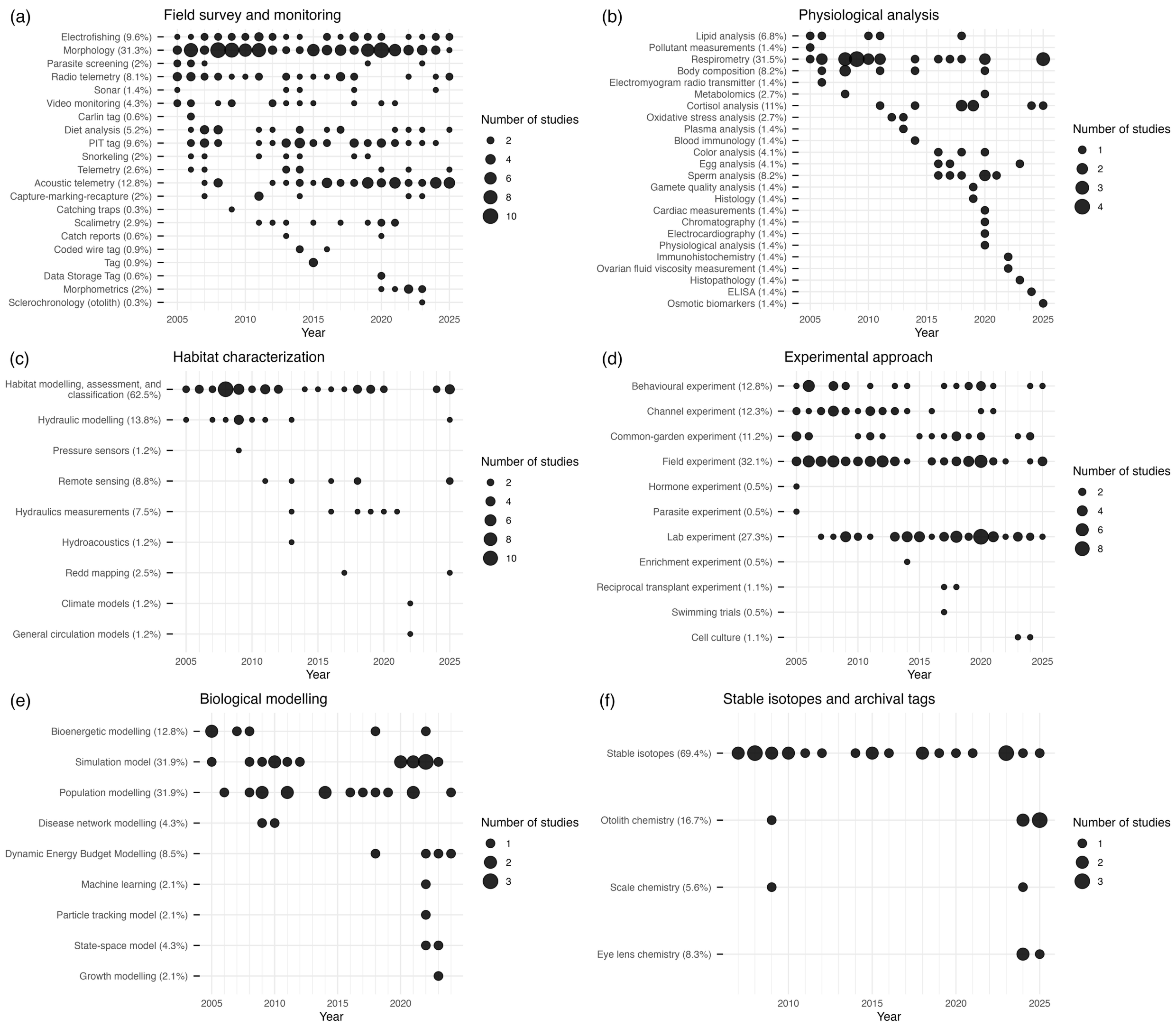


Figure S6: Temporal diversification of methods within (a) *Field survey and monitoring*, (b) *Physiological analyses*, (c) *Habitat characterization*, (d) *Experimental approach*, (e) *Biological modeling* and (f) *Stable isotopes and archival tags*.

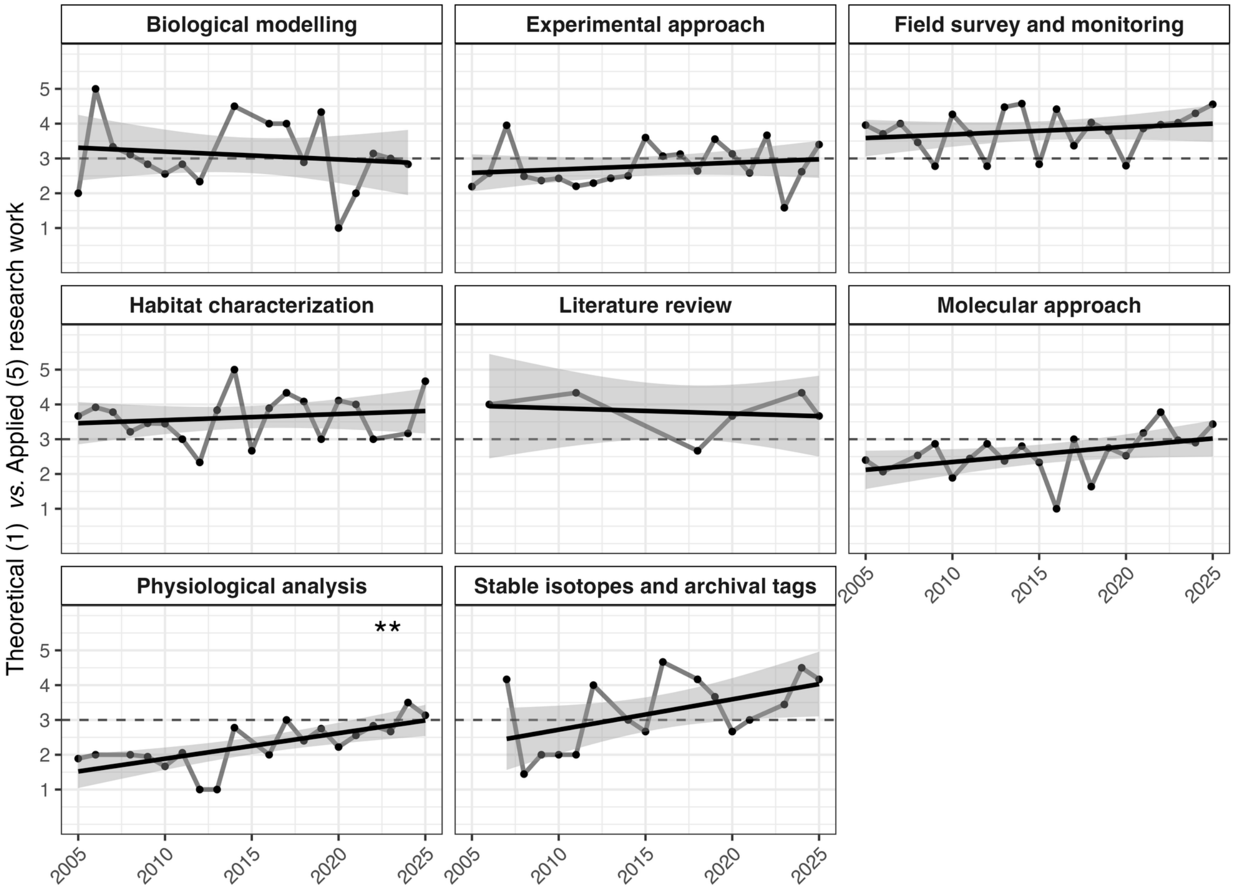


127

128 Figure S7: Temporal change of research positioning scores (theoretical: 1; applied: 5) according
129 to research methods. Points represent the annual average score obtained for each category.
130 Black solid lines show linear regression fits for each category, with shaded areas indicating 95%
131 confidence intervals around the fitted relationships. The dotted line represents a neutral value.
132 Asterisks denote statistically significant temporal trends based on the slope of the linear
133 regression ($p < 0.05$*, $p < 0.01$**, $p < 0.001$***).

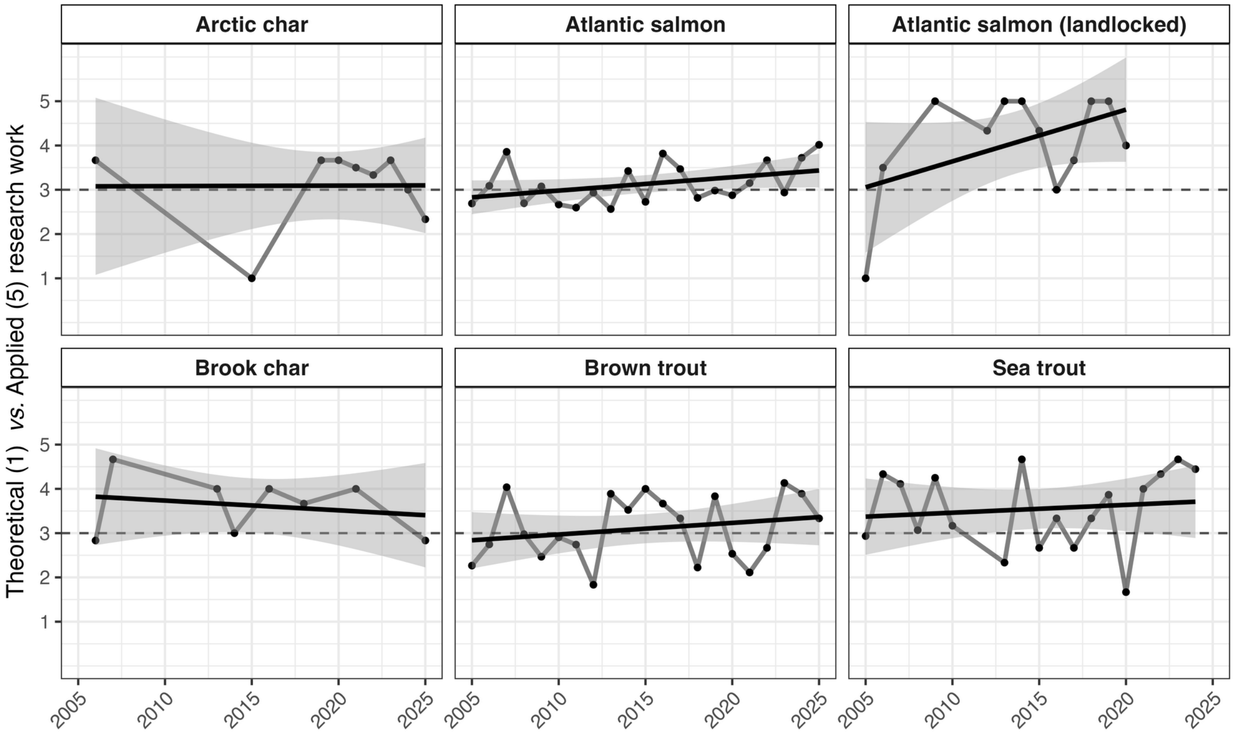


Figure S8: Temporal change of research positioning scores (theoretical: 1; applied: 5) according to study species. Points represent the annual average score obtained for each category. Black solid lines show linear regression fits for each category, with shaded areas indicating 95% confidence intervals around the fitted relationships. The dotted line reprents a neutral value. Asterisks denote statistically significant temporal trends based on the slope of the linear regression ($p < 0.05$*, $p < 0.01$**, $p < 0.001$***).

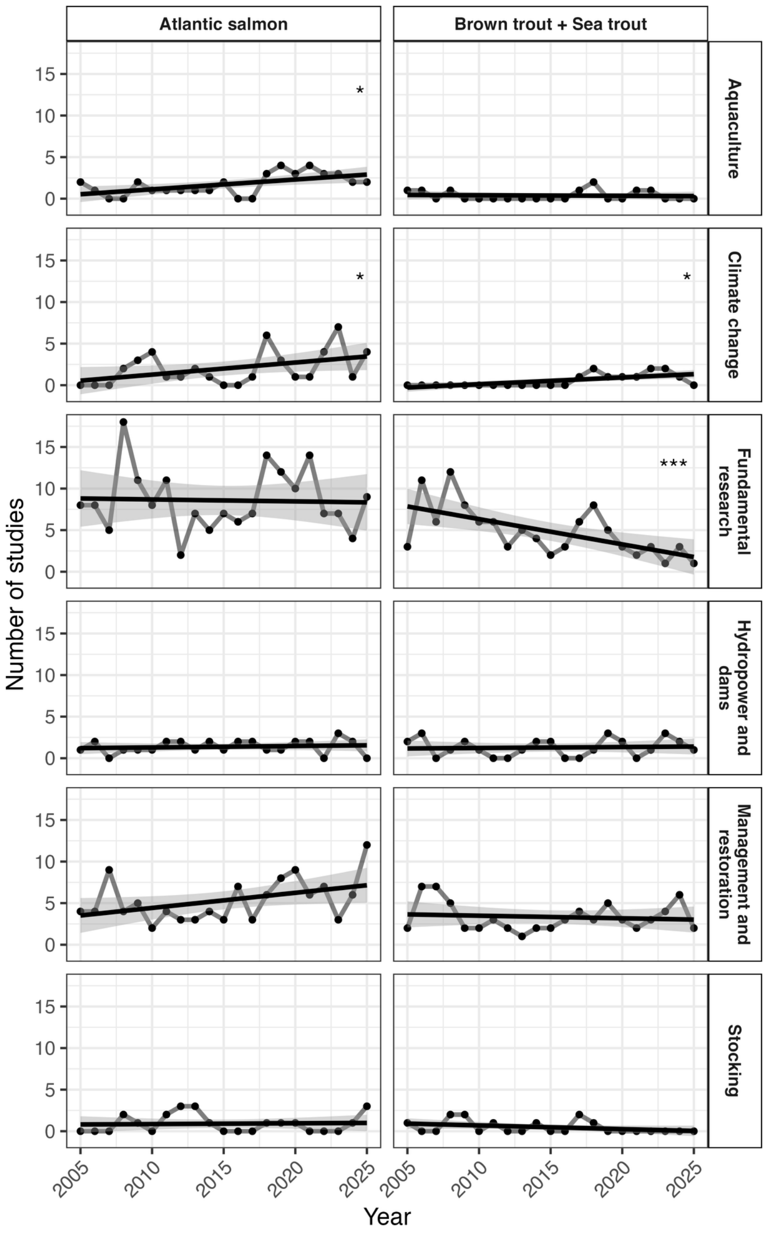


Figure S9: Temporal change in the occurrence of research drivers and for studies involving Atlantic salmon and brown trout and sea trout. Points represent the annual number of studies per category. Black solid lines show linear regression fits for each category, with shaded areas indicating 95% confidence intervals around the fitted relationships. Asterisks denote statistically significant temporal trends based on the slope of the linear regression ($p$ < 0.05*, $p$ < 0.01**, $p$ < 0.001***).

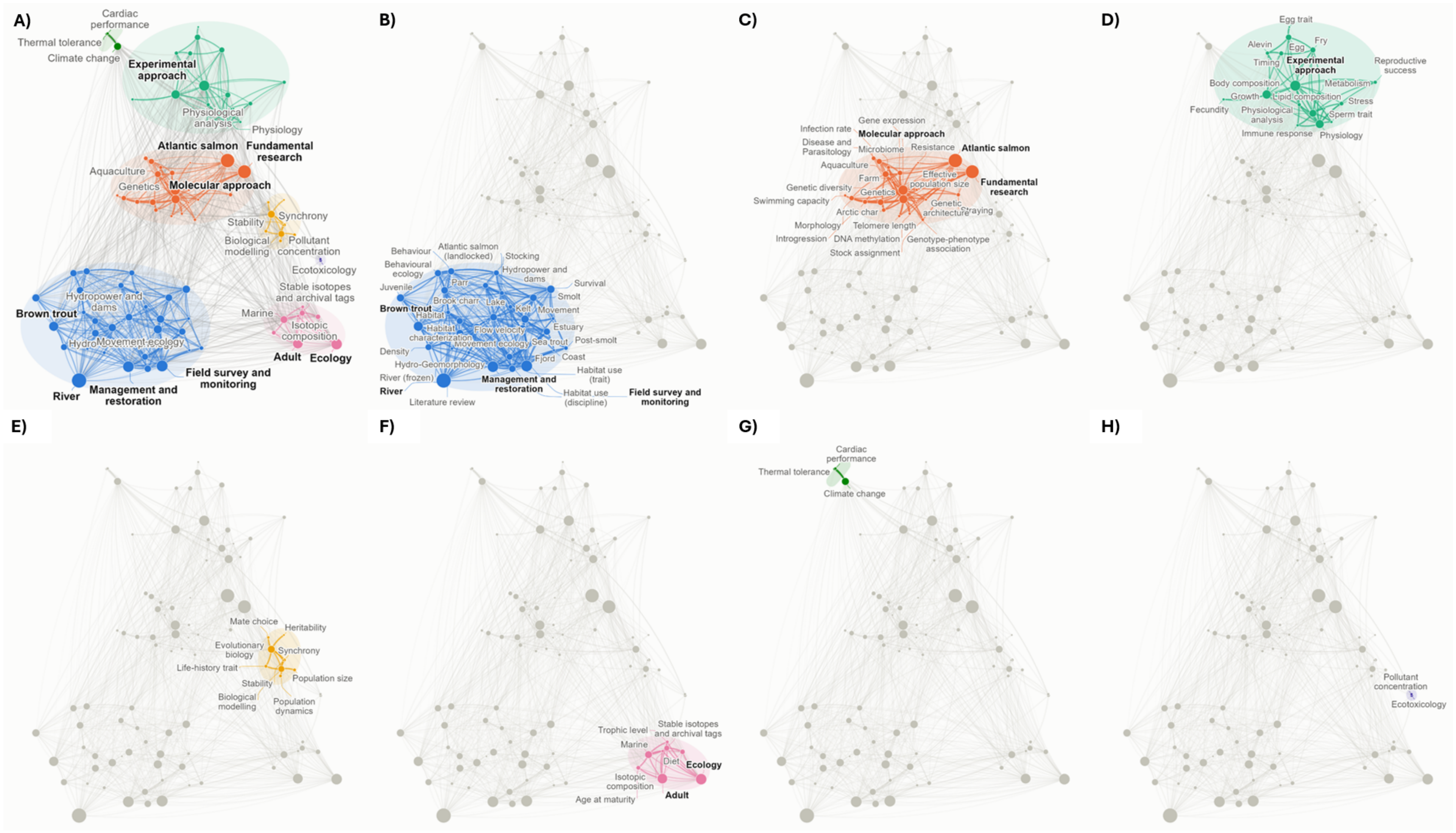


Figure S10: Full co-occurrence network (A) resulting from the 555 talks presented at the NoWPaS meetings between 2005 and 2025, and its seven clusters (B-H). Nodes are the categories of the seven coded variables (discipline, method, species, life stage, habitat, trait and driver), and node area is proportional to the number of talks featuring the category. Links connect pairs of categories cooccurring in at least three talks and are weighted by their association strength, *i.e.*, the ratio of observed to expected co-occurrences given the frequencies of the two categories (Van Eck and Waltman, 2010). Ellipses identify the seven clusters (C1–C7) obtained with the Louvain algorithm on the entire network (200 runs, Q = 0.42), identical to Figure 9.
